# Microtubule acetylation is a biomarker of cytoplasmic health during cellular senescence

**DOI:** 10.1101/2025.03.31.646469

**Authors:** Yusheng Shen, Roilea Maxson, Richard J. McKenney, Kassandra M. Ori-McKenney

**Affiliations:** Department of Molecular and Cellular Biology, University of California, Davis, Davis, CA 95616, USA

**Keywords:** Aging, microtubule acetylation, cytoplasm fluidity, cellular senescence

## Abstract

Cellular senescence is marked by cytoskeletal dysfunction, yet the role of microtubule post-translational modifications (PTMs) remains unclear. We demonstrate that microtubule acetylation increases during drug-induced senescence in human cells and during natural aging in *Drosophila*. Elevating acetylation via HDAC6 inhibition or *α*TAT1 overexpression in BEAS-2B cells disrupts anterograde Rab6A vesicle transport, but spares retrograde transport of Rab5 endosomes. Hyperacetylation results in slowed microtubule polymerization and decreased cytoplasmic fluidity, impeding diffusion of micron-sized condensates. These effects are distinct from enhanced detyrosination, and correlate with altered viscoelasticity and resistance to osmotic stress. Modulating cytoplasmic viscosity reciprocally perturbs microtubule dynamics, revealing bidirectional mechanical regulation. Senescent cells phenocopy hyperacetylated cells, exhibiting analogous effects on transport and microtubule polymerization. Our findings establish acetylation as a biomarker for cytoplasmic health and a potential driver of age-related cytoplasmic densification and organelle transport decline, linking microtubule PTMs to biomechanical feedback loops that exacerbate senescence. This work highlights the role of acetylation in bridging cytoskeletal changes to broader aging hallmarks.

## Introduction

Cellular aging is a gradual and irreversible process characterized by the accumulation of biochemical and biophysical changes that ultimately lead to cellular dysfunction (1). A central hallmark of aging is cellular senescence, a state of stable growth arrest that contributes to tissue decline and age-related diseases (2). While genetic, epigenetic, and metabolic factors driving aging and senescence have been extensively studied, the role of the cytoskeleton in these processes remains poorly understood (3–5). Aging is associated with increased protein aggregation (5–10) and declining energy metabolism (11, 12), both of which have been shown to disrupt intracellular transport (9, 13–15). Consistent with this, both impaired organelle transport (10, 16–19) and reduced microtubule dynamics (20) have been observed in aging cells, suggesting that changes in microtubule-based processes may be associated with cellular aging.

The microtubule cytoskeleton is dynamically regulated by microtubule-associated proteins (MAPs) and tubulin post-translational modifications (PTMs)(21–26). Key tubulin PTMs include acetylation of *α*K40 within the microtubule lumen and C-terminal tail modifications such as tyrosination/detyrosination and polyglutamylation, which influence microtubule stability, dynamics, and interactions with molecular motors (27–33). In neurons, aging is associated with the progressive accumulation of microtubule acetylation, detyrosination, and polyglutamylation, which are linked to microtubule stabilization and impaired axonal transport (17, 18, 24, 34, 35). These changes may contribute to age-related declines in organelle motility, synaptic function, and the accumulation of protein aggregates (6, 10). Interestingly, overall levels of protein acetylation rise significantly during muscle aging (36). However, why tubulin PTMs increase with age and how they directly influence microtubule dynamics and intracellular transport during aging or cell senescence have not been fully explored. It also remains unclear how age-related changes in PTMs interact with other hallmarks of aging, such as declining energy metabolism and altered cytoplasmic fluidity, to drive cellular dysfunction.

The cytoplasm is a densely packed, viscoelastic environment where macromolecules, organelles, and cytoskeletal networks interact to regulate intracellular processes (37, 38). Cytoplasmic fluidity, which governs the diffusion of molecules and organelles, is critical for maintaining cellular homeostasis and efficient intracellular transport (15, 32, 39, 40). During aging, the cytoplasm undergoes significant biophysical changes, including increased viscosity and reduced fluidity, which are associated with impaired organelle transport and protein aggregation (9, 10, 13). These changes are further exacerbated in senescent cells, where cytoplasmic stiffening has been linked to reduced metabolic activity (3, 41). Our previous work demonstrated that cells dynamically remodel their microtubule networks by modulating *α*-tubulin acetylation and MAP7 association in response to changes in cytoplasmic fluidity (32). The interplay between tubulin PTMs, cytoplasmic stiffening and cellular aging or senescence remains undefined. Defining these interactions could provide insights into the role of the cytoskeleton in aging and age-related diseases.

In this study, we investigate how the microtubule cytoskeleton is altered during cell senescence. We observed that increased microtubule acetylation is closely associated with induced cellular senescence *in vitro* and with aging *in vivo*. Using live-cell imaging and biochemical assays, we find that enhanced microtubule acetylation disrupts anterograde vesicle transport and increases pausing frequency during microtubule polymerization, effects not observed upon increasing detyrosination, highlighting specific roles for acetylation in altering these behaviors. We also explore the biophysical consequences of enhanced acetylation and find that it correlates with reduced cytoplasmic fluidity and impaired diffusion of micron-sized particles. Additionally, we show that modulating cytoplasmic viscosity independently affects microtubule polymerization dynamics, suggesting a reciprocal relationship between the cytoskeleton and cytoplasmic rheology. Strikingly, senescent cells exhibit intracellular transport dynamics similar to those of hyperacetylated cells, reinforcing the idea that microtubule acetylation plays a mechanistic role in aging-associated transport defects. Collectively, our findings establish direct links between acetylation, cytoplasmic mechanics, and intracellular transport dysfunction, providing mechanistic insight into how cytoskeletal modifications contribute to cellular senescence.

## Results

### Microtubule acetylation is associated with cell senescence *in vitro* and aging *in vivo*

To study how cells optimize their microtubule cytoskeletons during the state of cell senescence, we treated BEAS-2B cells, a human lung epithelial cell line, with Doxorubicin (Doxo), a DNA damaging agent commonly used to induce cell senescence *in vitro* (41–43). We observed a sharp decrease (87.9%) of 5-bromo-2-deoxyuridine (BrdU) incorporation into newly synthesized DNA and 1.4-fold increase in cell projection area in Doxo-treated cells (Fig. S1A-E), consistent with the growth-arrest features reported for senescent cells both *in vitro* and *in vivo* (41, 42). We then used immunostaining to analyze microtubule acetylation, detyrosination, and polyglutamylation. Surprisingly, we observed a marked increase in microtubule acetylation (4.1-fold) accompanied by a 1.4-fold increase in microtubule mass, as well as a 1.4-fold increase in detyrosination, while polyglutamylation levels decreased by a third (Figs. 1B-D). Because we observed the most pronounced effect on acetylation following Doxo treatment, we examined whether increased microtubule acetylation was a universal feature of drug-induced cell senescence. We treated BEAS-2B cells with other senescence-inducing drugs, Camptocethin (CPT) and Cisplatin (44), which also displayed growth arrest features (Fig. S1A-E). Remarkably, we observed similar increases in acetylation and microtubule mass for both CPT- and Cisplatin-treated cells (Figs. 1E-H). Our results coupled with previous studies using different drugs to induce cell senescence (45, 46) suggest that increased microtubule acetylation is coupled with cell senescence *in vitro*.

**Fig. 1.**
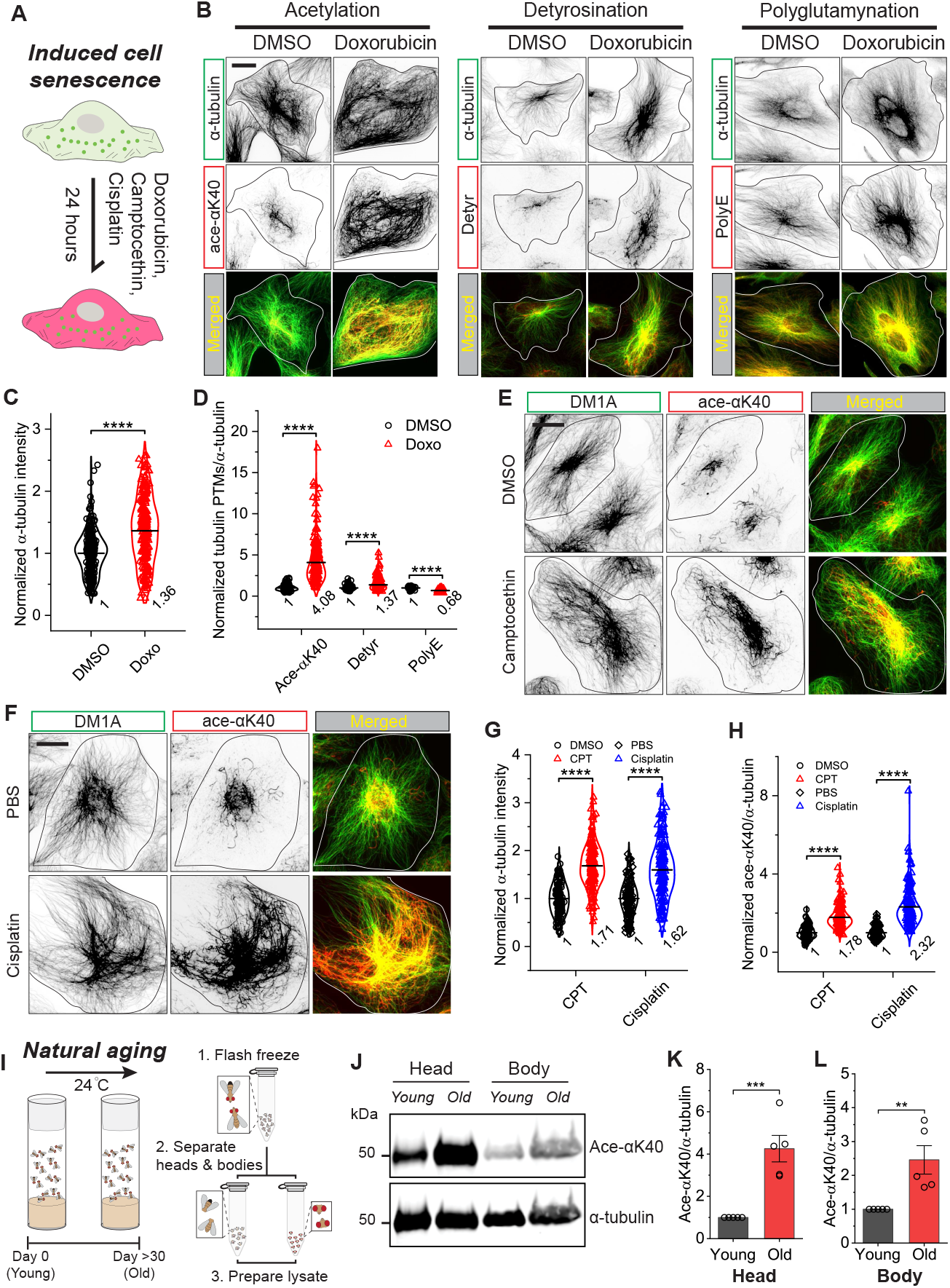
Microtubule acetylation is associated with aging both *in vitro* and *in vivo*. (A) Schematic illustration of induced cell senescence by treating cells with Doxorubicin (Doxo), Camptocethin (CPT) or Cisplatin for 24 hours. (B-D) Representative images and quantification comparing *α*-tubulin intensity (C) and relative changes of tubulin PTMs (D) for DMSO (control) or 200 ng/mL Doxorubicin-treated (Doxo) cells. Acetylation at Lys40 (Ace-*α*K40), detyrosination (Detyr) and polyglutamylation (PolyE) were assessed relative to the total *α*-tubulin. Ace-*α*K40, n = 282 and 235 cells from 3 independent experiments, \*\*\*\**p <* 0.0001; Detyr, n = 189 and 207 cells from 3 independent experiments, \*\*\*\**p <* 0.0001; PolyE, n = 100 and 86 cells from 3 independent experiments, \*\*\*\**p <* 0.0001. Scale bar: 20 *µ*m. (E-H) Representative images and quantification comparing *α*-tubulin intensity (G) and Ace-*α*K40/*α*-tubulin ratios (H) for DMSO/PBS (control) or Camptocethin (CPT, 200 nM) or Cisplatin (50 *µ*M) treated cells. CPT, n = 138 and 172 cells from 3 independent experiments, \*\*\*\**p <* 0.0001; Cisplatin, n = 123 and 176 cells from 3 independent experiments, \*\*\*\**p <* 0.0001. Scale bar: 20 *µ*m. (I) Schematic illustration of the procedures for preparing young (0 DAE) and old (30-40 DAE) *Drosophila* samples for assessing the microtubule acetylation. (J-L) Western-blots (WB) and corresponding quantification showing relative changes of acetylated *α*-tubulin in the head and body for young and old *Drosophila*. n = 5 independent experiments, \*\*\**p* = 0.00084, \*\**p* = 0.0084. All violin graphs display all data points with means. P-values were calculated using a student’s t-test.

To investigate whether microtubule acetylation also increases during natural aging *in vivo*, we utilized *Drosophila melanogaster*. This model organism exhibits many age-related functional changes observed in humans, such as alterations in motor activity, learning and memory, cardiac function, and fertility (47). We allowed male and female flies to age naturally at 24 ^°^C and collected adult flies at two time points: 0 days after eclosion (DAE, young) and 30-40 DAE (old). Western blot analysis on lysates made from separated heads and bodies (Fig. 1I) revealed significant increases in *α*-tubulin acetylation levels in both the heads (4.2-fold) and the bodies (2.3-fold) of old flies compared to young flies (Figs. 1J-L). Acetylation levels were generally much higher in the brain than in the body (Fig. 1J), consistent with microtubule acetylation enrichment in neuronal cells (47–49). These findings demonstrate that acetylation increases with age in *Drosophila*.

### Elevated cellular acetylation disrupts anterograde vesicle transport

To understand why cells might promote microtubule acetylation during senescence, we sought to characterize the effects of enhanced microtubule acetylation in BEAS2B cells. To manipulate microtubule acetylation, we used two approaches: pharmacological inhibition of histone deacetylase-6 (HDAC6) with tubacin, a widely used drug that increases microtubule acetylation (32, 50–55), and transient expression of wild type or catalytically dead (D157N) *α*-tubulin acetyl transferase-1 (*α*TAT1: acetylating enzyme) (Figs. 2A and E) (56, 57). Similar to prior results (32), tubacin treatment induced a 10.5-fold increase in acetylation without affecting microtubule mass or organization (Figs. 2B–D). Similarly, wild-type *α*TAT1 expression increased acetylation by 15.8-fold but reduced microtubule mass by 18.2% compared to the D157N control (Figs. 2F–H). Both methods efficiently elevated microtubule acetylation in BEAS-2B cells, enabling us to study its effects on a range of microtubule-based processes.

**Fig. 2.**
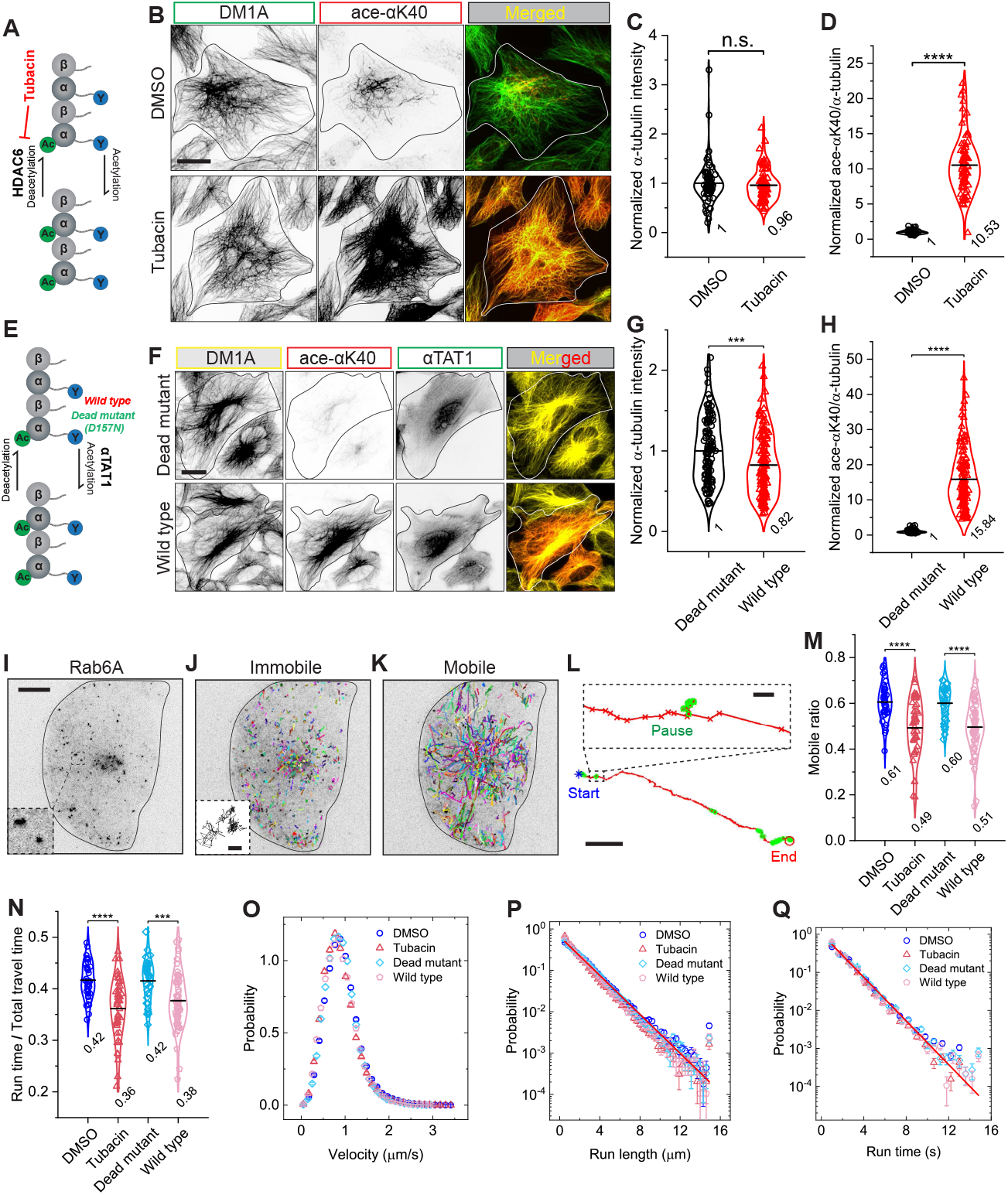
Microtubule acetylation impacts anterograde vesicle transport. (A-D) Representative images and quantification comparing *α*-tubulin intensity (C) and acetylation at Lys40 (Ace-*α*K40)/*α*-tubulin ratios (D) for DMSO (control) or 5 *µ*M tubacin-treated cells. n = 72 and 81 cells, respectively, from 3 independent experiments, \*\*\*\**p <* 0.0001. Scale bar: 20 *µ*m. (E-G) Representative images and quantification comparing *α*-tubulin intensity (G) and Ace-*α*K40/*α*-tubulin ratios (H) for cells transiently expressing the wild type (wild type) or catalytically dead mutant (D157N, dead mutant) of *α*TAT1-mScarlet. n = 120 and 129 cells, respectively, from 3 independent experiments, \*\*\**p* = 0.00044, \*\*\*\**p <* 0.0001. Scale bar: 20 *µ*m. (I) Image of a BEAS-2B cell expressing EGFP-Rab6A visualized by TIRF microscopy. Inset: magnified view of Rab6A vesicles. Scale bar, 10 *µ*m. (J and K) Representative trajectories (colored lines) of EGFP-Rab6A-positive secretory vesicles (10 fps for 3 min) showing the relative immobile (J) and mobile (K) fractions. Inset shows a magnified view of an immobile vesicle trajectory. Scale bar: 100 nm. (L) Magnified trajectory image showing a Rab6A-positive vesicle. Blue stars and red circles mark the start and end point of the trajectory, respectively. Pause states are highlighted by green circles. Motion states were detected and extracted by a homemade algorithm (32, 58, 68). Scale bar: 2 *µ*m. Inset shows a magnified view of the pause state. Scale bar: 100 nm. (M and N) Measured mobile ratio and run time/total travel time of Rab6A-positive vesicles in response to increased microtubule acetylation induced by either treating cells with DMSO/tubacin or transiently expressing the dead mutant/wild type *α*TAT1. \*\*\**p* = 0.00039, \*\*\*\**p <* 0.0001. Each data point represents the average of measured trajectories from a single cell. (O-Q) Measured probability density functions (PDFs) of trajectory-based run velocity, *v*, run length, *L*_*p*_, and run time, *T*_*p*_, for Rab6A-vesicles in DMSO or tubacin-treated cells, or in cells transiently expressing expressing the wild type (wild type) or catalytically dead mutant (D157N, dead mutant) of *α*TAT1. The measured peak velocities for DMSO and tubacin treated cells, and for cells expressing dead mutant and wild type *α*TAT1 are 0.89 ± 0.02, 0.77 ± 0.02, 0.85 ± 0.02 and 0.83 ± 0.02 *µ*m/s, respectively. The measured *L*_*p*_ and *T*_*p*_ both could be fitted to a simple exponential distribution (red solid lines, only the fittings to the DMSO-treated cells were shown for simplicity), with a characteristic *L*_*p*_ = 1.85 ± 0.02, 1.52 ± 0.02, 1.81 ± 0.02 and 1.59 ± 0.02 *µ*m, and *T*_*p*_ = 1.60 ± 0.04, 1.37 ± 0.02, 1.51 ± 0.04 and 1.35 ± 0.04 s, respectively, for DMSO or tubacin-treated cells and for cells expressing dead mutant or wild type *α*TAT1. Total trajectories analyzed from the mobile fraction are n = 37926, 32609, 30867 and 31820 from n = 43, 50, 35 and 58 cells from more than 3 independent experiments. All violin graphs display all data points with means. P-values were calculated using a student’s t-test.

First, we investigated the effects of elevated microtubule acetylation on anterograde and retrograde vesicle transport. We conducted a comprehensive analysis of vesicle motion dynamics in living cells (32, 58). While previous studies have documented preferential binding of kinesin-1 to acetylated microtubules (52, 59–61), the impact of acetylation on kinesin-driven transport of micron-sized vesicle cargoes in particular remains unclear. These vesicles navigate a mechanically and biochemically complex cytoplasm, encountering impedance from poro- and visco-elastic properties (15, 39, 40, 62) and extensive biochemical interactions (63–66). We focused on the transport of secretory vesicles (EGFP-Rab6A) primarily driven anterogradely by kinesin-1 and kinesin-3 motors (32, 67). Rab6A-vesicle motion exhibited significant heterogeneity, consisting of immobile tracks without directed motion and mobile tracks with persistent, directed motion (Figs. 2I–K). Motile vesicles displayed a two-state behavior, switching between diffusive or paused states and directed runs (Fig. 2L), arising from interactions between the motor-cargo and the cytoplasmic environment (32, 58). To characterize this behavior, we applied a previously developed algorithm (32, 58, 68) to extract distinct motion states. We analyzed the mobile fraction of vesicles, the ratio of run time to total travel time for mobile vesicles, as well as the velocities, run lengths, and run times of individual transport segments, providing a detailed and unbiased assessment of the motion dynamics.

Both tubacin treatment and *α*TAT1 expression resulted in similar reductions in vesicle mobility, with mobile ratios decreasing by 19.7% and 15.0%, respectively, and run time/total travel time ratios decreasing by 14.3% and 9.5% compared to controls (Figs. 2M and N). Probability density functions (PDFs) of run velocities exhibited similar peaks, ranging from 0.77 to 0.89 *µ*m/s, with tubacin treatment and *α*TAT1 expression causing a 13.5% and 2.4% decrease in peak velocity (Fig. 2O). PDFs of run length and run time followed exponential distributions, with characteristic values of 1.52–1.85 *µ*m for run length and 1.35–1.60 s for run time (Figs. 2P and Q). Elevated acetylation reduced run length by 17.8% and 12.2% and run time by 14.4% and 10.6% for tubacin and *α*TAT1 expression, respectively. However, the same analysis performed on retrogradely transported Rab5-positive early endosomes found that all 5 parameters showed no significant changes upon tubacin treatment (Fig. S2). These findings indicate that elevated microtubule acetylation differentially affects anterograde and retrograde motor-driven transport dynamics by negatively impacting anterograde cargo trafficking.

### Microtubule acetylation, but not detyrosination, induces frequent pausing during microtubule growth

We next wanted to examine the effects of increased acetylation on microtubule polymerization. By expressing the +TIP marker, EB1-tdTomato, in BEAS-2B cells, we observed comet-like structures growing from the microtubule organizing center (MTOC) to the cell edge as expected (69) (Fig. 3A and Movie S1). Using single particle tracking (SPT) (32, 70), we measured mean-squared displacements (MSDs) for comet trajectories, from which we computed the polymerization rate and polymerization length of each trajectory (Figs. 3B-D). The measured PDFs of polymerization rates in DMSO and tubacin-treated cells both featured a Gaussian distribution (Fig. 3F), which agrees with previous *in vivo* findings (69, 71, 72). Tubacin treatment reduced polymerization rate by 37.0%, polymerization length by 24.8%, and the total number of polymerization events by 50.2% (Figs. 3E–H).

**Fig. 3.**
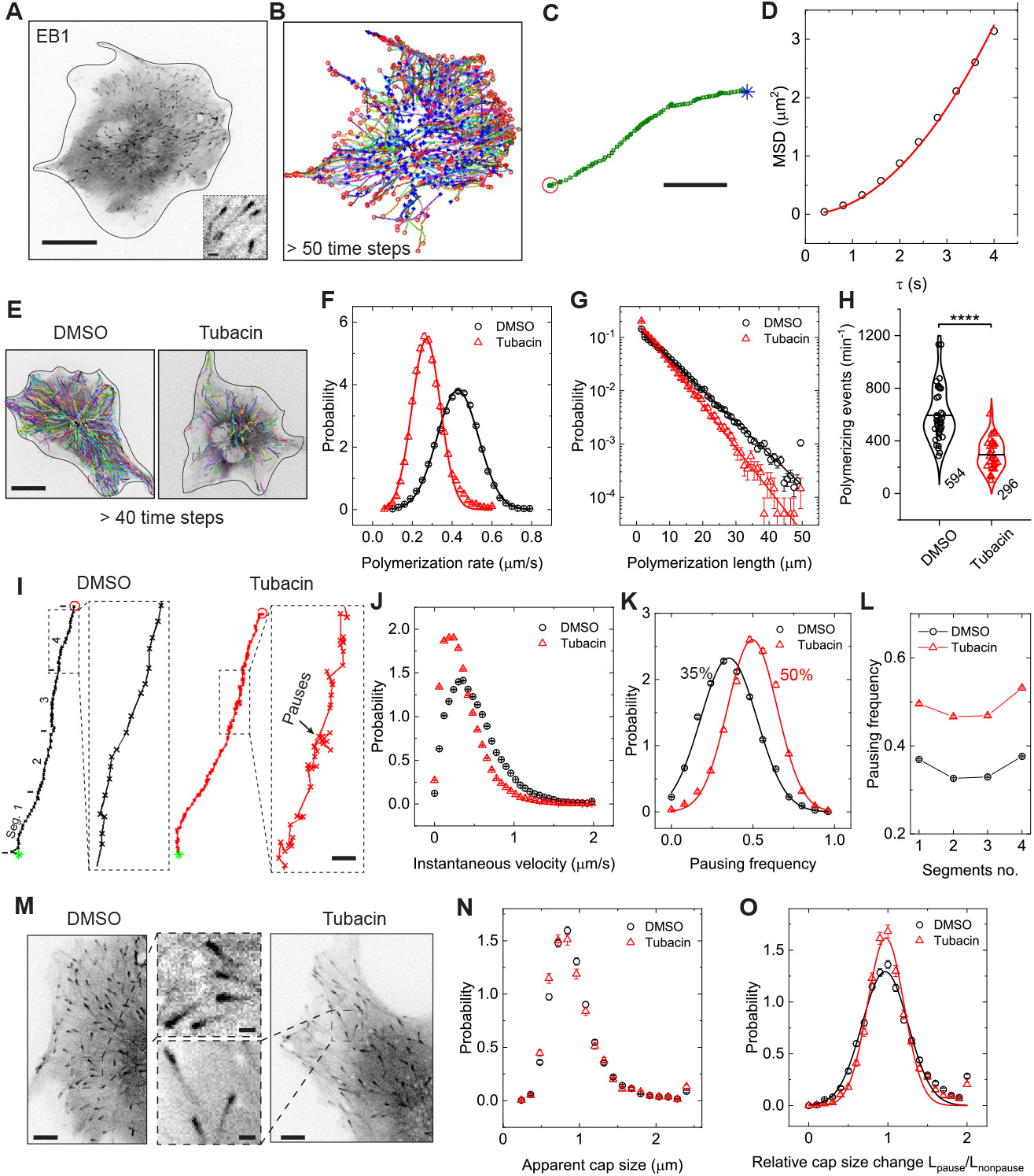
Tubacin treatment decreases growing microtubule polymerization rates by inducing frequent pausing. (A) Image of a BEAS-2B cell expressing EB1-tdTomato visualized by spinning disk confocal microscopy. Scale bar: 20 *µ*m. Inset, magnified view of EB1 comets. Scale bar: 1 *µ*m. (B) Representative trajectories (colored lines) of EB1 comets (2.5 fps for 3 min) showing the growth pattern of microtubules in cells. Blue stars and red circles mark the start and end point of each trajectory, respectively. (C) A representative EB1 trajectory. Scale bar: 5 *µ*m. (D) Measured MSD as a function of delay time *τ* for a representative EB1 trajectory as shown in (C). (E) Representative trajectories of EB1 comets for DMSO or tubacin treated cells. Scale bar, 20 *µ*m. (F and G) Measured PDFs of the polymerization rate (F) and the polymerization length (G) for individual EB1 comets in DMSO (black circles) or tubacin treated (red triangles) cells. The solid lines in (F) shows the fitting of the data points to Gaussian distribution with the mean polymerization rates are 0.43 ± 0.01 and 0.27 ± 0.01 *µ*m/s, respectively. The measured polymerization length in (G) could be fitted to a simple exponential distribution (solid lines) with a characteristic length = 7.3 ± 0.14 and 5.5 ± 0.12 *µ*m, respectively, for DMSO or tubacin-treated cells. (H) Measured polymerizing events per minute in DMSO or tubacin-treated cells. \*\*\*\**p <* 0.0001. Total EB1 trajectories analyzed are n = 58798 and 20643 from n = 38 and 34 cells from 3 independent experiments. (I) Magnified trajectory images showing EB1-comets movements in DMSO or tubacin-treated cells. Green stars and red circles mark the start and end point of the trajectory, respectively. Scale bar, 0.5 *µ*m. (J and K) Measured PDFs of the instantaneous velocity of EB1-comets (J) and pausing frequency (K) for DMSO or tubacin-treated cells. The measured peak velocities for DMSO and tubacin treated cells are 0.36 ± 0.05 and 0.20 ± 0.05 *µ*m/s, respectively. The solid lines in (K) show the fitting of the data points to Gaussian distribution with the mean pausing frequencies are 35 ± 0.4 % and 50 ± 0.5 %, respectively. (L) Measured pausing frequency as a function of the trajectory-segment no. for DMSO or tubacin-treated cells. To quantify the temporal dependence of pausing frequency, individual trajectories (*>* 20 time steps) were cut into 4 segments as shown in (I) with segment 1 being the initiating phase and 4 being the ending phase of a polymerization event, respectively. (M) Images of BEAS-2B cells expressing EB1-tdTomato and treated with either DMSO or tubacin. Scale bar: 5 *µ*m. Cropped images, magnified view of EB1 comets. Scale bar: 1 *µ*m. (N) Measured PDFs of the apparent EB1-comet size in cells treated with either DMSO or tubacin. (O) Measured PDFs of the relative changes of EB1-comet size during the pausing or non-pausing states of individual trajectories in cells treated with either DMSO or tubacin. Total EB1 trajectories analyzed are n = 58798 and 20643 from n = 38 and 34 cells from 3 independent experiments. All violin graphs display all data points with means. P-values were calculated using a student’s t-test.

Microtubule acetylation and detyrosination are typically associated with “stable microtubules” (24, 27, 73) and often occur on the same subset of microtubules (73–76). We were therefore curious to examine whether microtubule “stabilization” by detyrosination could exert similar effects on microtubule polymerization. Detyrosination of *α*-tubulin is catalyzed by an enzyme complex composed of a vasohibin (VASH1 or VASH2) and a small vasohibin-binding protein (SVBP) (77, 78). To manipulate detyrosination in cells, we transiently expressed the wild type or catalytic dead mutant (C168A) of the enzyme in BEAS-2B cells (Fig. S3A). Immunostaining analysis revealed the detyrosination signal increased 5.0-fold and expanded to nearly all microtubules in the wild-type-expressing cells compared to control cells, whereas the overall microtubule intensity remained unchanged (Figs. S3B-D). VASH1 expression slightly increased polymerizing events 1.3-fold (Figs. S3E-H) compared to controls. Thus, acetylation, but not detyrosination, impedes microtubule polymerization in cells by reducing growth rate, increasing catastrophe frequency, and decreasing the total number of polymerizing microtubules. Interestingly, these polymerization parameters are unaffected by acetylation or detyrosination in *in vitro* assays with purified proteins (30, 35), suggesting the effects we observe arise from an altered cellular environment.

In cells, the apparent polymerization rate or mobility of microtubule plus-ends is governed by the rate of GTP-tubulin dimer addition and the frequency of dynamic pausing states (38, 79–81). To gain a better understanding of how microtubule acetylation affects polymerization dynamics, we analyzed the instantaneous velocity and pausing behaviors of individual EB1 comet trajectories (Figs. 3I-L). Tubacin treatment slowed EB1 comets and increased pausing without inducing catastrophe, as evidenced by shorter single-moving steps and direction reversals (Fig. 3I and Movie S2). The PDF of instantaneous velocity showed a 44.4% reduction in tubacin-treated cells, indicating a decreased rate of GTP-tubulin addition (Fig. 3J). The PDF of pausing frequency displayed a Gaussian distribution, with a 1.4-fold increase in tubacin-treated cells (Fig. 3K). Notably, we observed a 35.0% basal pausing frequency in control cells, suggesting dynamic pausing is a common feature of growing microtubules. Further temporal analysis of EB1 trajectories revealed that pausing frequency was consistent throughout polymerization, with slight increases at initiation and termination phases (Fig. 3L). This suggests pausing occurs randomly, with no temporal or spatial preference even after acetylation. Cells expressing *α*TAT1 also showed inhibited microtubule growth (Fig. S4). In contrast, VASH1 expression did not alter instantaneous velocity or pausing frequency compared to controls (Figs. S3I-L).

In cells, the apparent microtubule growth rate has been shown to positively correlate with the GTP-cap size (82–84). We next examined whether the observed changes in polymerization rate and pausing behavior in hyperacetylated cells are associated with changes in GTP-cap size. However, the apparent EB1-comet size remained unchanged in tubacin-treated cells compared to controls (Figs. 3M and N). Furthermore, we observed no distinguishable changes in comet size when the growing microtubule plus-end was either in the pausing or non-pausing state (Fig. 3O). Overall, we find that hyper-acetylation perturbs nascent microtubule growth by reducing instantaneous velocity and increasing pausing at the plus ends independently of GTP-cap size.

### Modulating cytoplasm fluidity affects microtubule polymerization dynamics

How could increased acetylation alter the growth of microtubules in cells considering this modification does not alter microtubule dynamics in reconstitution studies with purified proteins (35)? Previous studies have demonstrated that physical cues such as cytoplasm fluidity are involved in the regulation of microtubule polymerization (38, 85). We therefore sought to understand how cytoplasm fluidity affects EB1 comet dynamics by using both physical and biochemical approaches. We either treated cells with a hypertonic solution to increase cytoplasm density or with blebbistatin, which inhibits myosin II motor activity. Both approaches have been shown to effectively decrease cytoplasm fluidity (14, 32, 38, 58, 86). Interestingly, both treatments decreased polymerization rate (18.2% and 26.1%, respectively), polymerization length (8.2% and 17.6%, respectively), and number of polymerization events (20.3% and 37.7%, respectively) (Figs. 4A-H). Decreases in polymerization rates were also accompanied by decreases in instantaneous velocities (Figs. 4I and J) and increases in pausing behaviors (Figs. 4K-M). Therefore, modulating cytoplasm fluidity caused similar changes in the dynamic behaviors of growing microtubules as observed in hyper-acetylated cells.

**Fig. 4.**
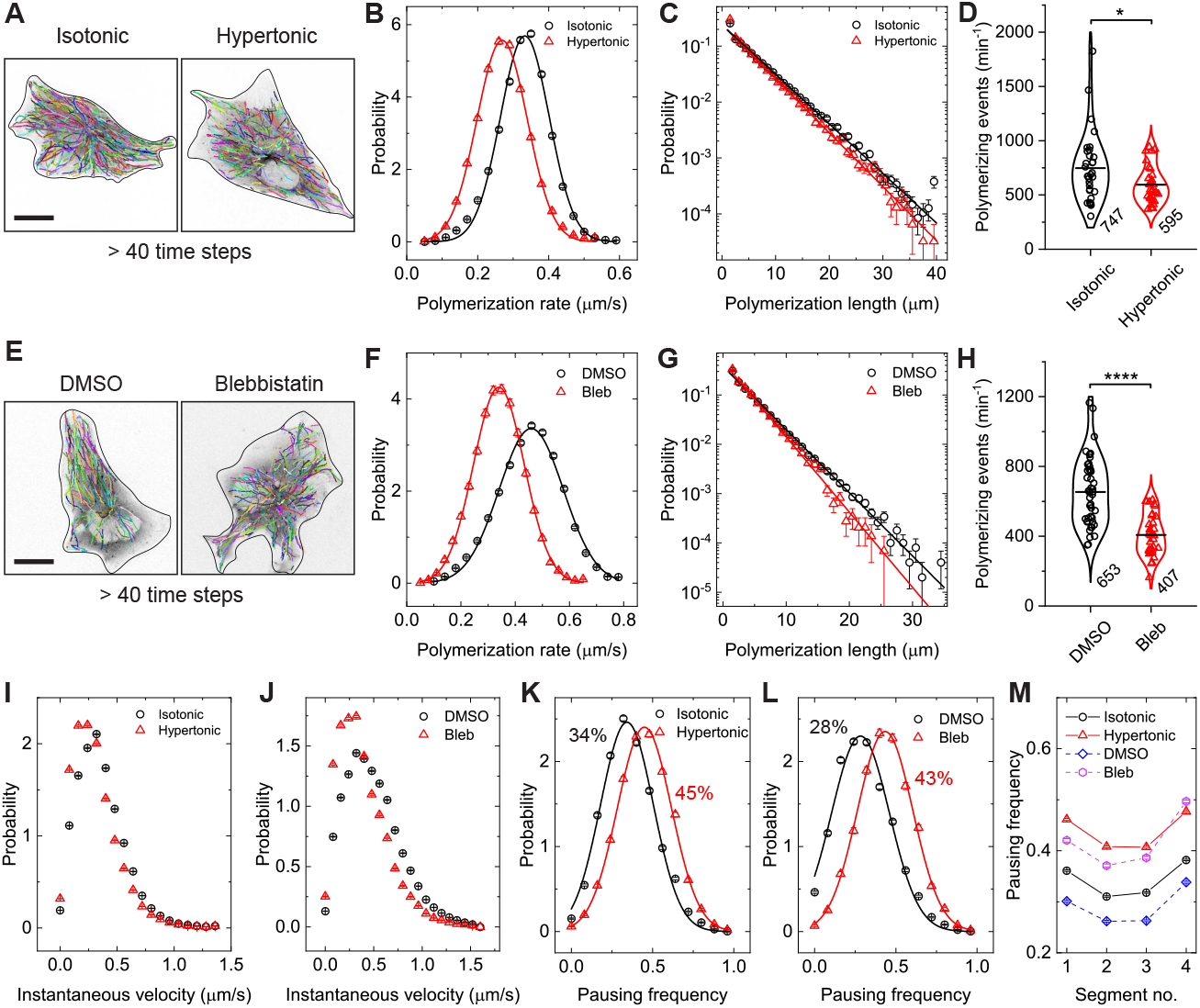
Modulating cytoplasm fluidity affects microtubule polymerization dynamics. (A) Representative trajectories of EB1 comets in cells treated with isotonic (310 mOsm) or hypertonic solutions (400 mOsm). Scale bar, 20 *µ*m. (B and C) Measured PDFs of the polymerization rate (B) and the polymerization length (C) for individual EB1 comets in cells treated with either isotonic or hypertonic solutions. The solid lines in (B) shows the fitting of the data points to Gaussian distribution with the mean polymerization rates are 0.33 ± 0.01 and 0.27 ± 0.01 *µ*m/s, respectively. The measured polymerization length in (C) could be fitted to a simple exponential distribution (solid lines) with a characteristic length = 4.9 ± 0.15 and 4.5 ± 0.15 *µ*m, respectively, for cells treated with either isotonic or hypertonic solutions. (D) Measured polymerizing events per minute in cells treated with either isotonic or hypertonic solutions. \**p* = 0.022. Total EB1 trajectories analyzed are n = 47773 and 31088 from n = 32 and 31 cells from 3 independent experiments. (E) Representative trajectories of EB1 comets in DMSO- or blebbistatin-treated cells. Scale bar, 20 *µ*m. (F and G) Measured PDFs of the polymerization rate (F) and the polymerization length (G) for individual EB1 comets in DMSO- or blebbistatin-treated (Bleb) cells. The solid lines in (F) shows the fitting of the data points to Gaussian distribution with the mean polymerization rates are 0.46 ± 0.01 and 0.34 ± 0.01 *µ*m/s, respectively. The measured polymerization length in (G) could be fitted to a simple exponential distribution (solid lines) with a characteristic length = 3.4 ± 0.15 and 2.8 ± 0.15 *µ*m, respectively, in DMSO- or blebbistatin-treated cells. (H) Measured polymerizing events per minute in DMSO- or blebbistatin-treated cells. \*\*\*\**p <* 0.0001. Total EB1 trajectories analyzed are n = 49794 and 14647 from n = 44 and 35 cells from 3 independent experiments. (I and J) Measured PDFs of the instantaneous velocity of EB1-comets in cells treated with isotonic or hypertonic solutions (I), and in DMSO- or blebbistatin-treated cells (J). The measured peak velocities are 0.32 ± 0.05, 0.20 ± 0.05, 0.35 ± 0.05 and 0.25 ± 0.05 *µ*m/s, respectively. (K and L) Measured PDFs of pausing frequency for individual EB1 comets in cells treated with isotonic or hypertonic solutions (K), and in DMSO- or blebbistatin-treated cells (L). The solid lines in (K and L) show the fitting of the data points to Gaussian distribution with the mean pausing frequencies are 34 ± 0.5 % and 45 ± 0.2 % for cells treated with isotonic or hypertonic solutions, respectively, and 28 ± 0.6 % and 43 ± 0.2 % in DMSO- or blebbistatin-treated cells, respectively. (M) Measured pausing frequency as a function of the trajectory-segment no. for cells treated with isotonic or hypertonic solutions (open symbols connected with solid lines), and in in DMSO- or blebbistatin-treated cells (open symbols connected with dashed lines). All violin graphs display all data points with means. P-values were calculated using a student’s t-test.

### Microtubule acetylation decreases cytoplasmic fluidity and impedes micron-sized particle diffusion

Based on the similar effects on microtubule dynamics that we observed upon increased acetylation and increased cytoplasmic density, we investigated how enhanced acetylation affects cytoplasmic fluidity. To measure changes in fluidity, we performed micro-rheology in BEAS-2B cells using 40-nm diameter genetically encoded multimeric nanoparticles (40-nm GEM) as tracer particles (Fig. 5A) (32, 87). These GEMs are comparable in size to ribosomes and other macro-molecular complexes in cells (87–89). Using single-particle tracking (32, 70), we tracked GEM diffusion and calculated the MSDs for individual particles. The MSDs exhibited a linear relationship with delay time (*τ*), indicating free diffusion of GEMs in BEAS-2B cells (Fig. 5B). Interestingly, while tubacin treatment did not alter GEM diffusion, VASH1 expression increased diffusion by 1.2-fold (Figs. 5C and D). We also confirmed that blebbistatin treatment caused a 48.4% decrease of diffusion of GEMs, indicating decreased cytoplasm fluidity (Fig. S5). These results suggest that the inhibition of microtubule polymerization caused by acetylation is not due to reduced diffusion or a slower addition rate of GTP-tubulin dimers to microtubule plus-ends (38).

**Fig. 5.**
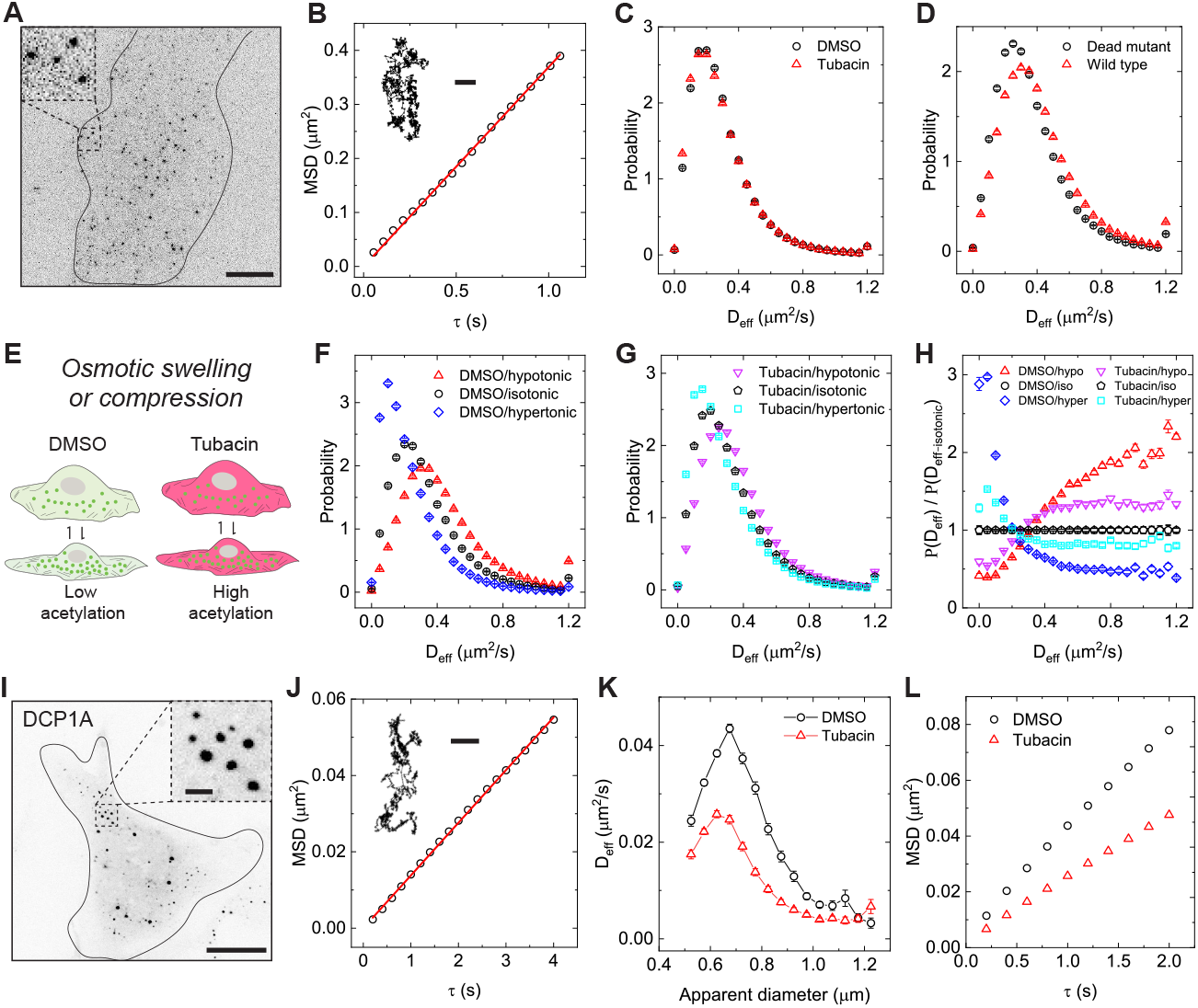
Microtubule acetylation and detyrosination differentially tune cytoplasm fluidity. (A) Image of a BEAS-2B cell expressing 40-nm GEMs visualized by TIRF microscopy. Inset: magnified view of GEMs. Scale bar, 10 *µ*m. (B) Measured MSD as a function of delay time *τ* for a representative GEMs trajectory as shown in the inset. Scale bar: 1 *µ*m. (C and D) Measured PDFs of the effective diffusion coefficient *D*_eff_ for individual GEMs in DMSO or tubacin treated cells (C) and for GEMs in cells transiently expressing the wild type or catalytically dead mutant of VASH1 (D). The measured peak *D*_eff_ are 0.18 ± 0.01, 0.18 ± 0.01, 0.31 ± 0.01 and 0.25 ± 0.01 *µ*m^2^/s, respectively. Total GEMs trajectories analyzed are n = 197620 and 277040 from n = 57 and 53 cells from 4 independent experiments for acetylation experiments and n = 138981 and 139802 from n = 57 and 63 cells from 4 independent experiments for detyrosination experiments. (E) Schematic illustration showing the measurements of the diffusion dynamics of GEMs in cells with either low microtubule acetylation (DMSO treated) or high acetylation (tubacin treated) and then subjected to different osmotic treatments to either compress or swell the cell. (F and G) Measured PDFs of *D*_eff_ for GEMs in cells with either low acetylation (DMSO treated, F) or high acetylation (tubacin treated, G) and then subjected to different osmotic treatments (isotonic, 310 mOsm; hypotonic, 250 mOsm; hypertonic, 400 mOsm). (H) Comparison of the relative change *P* (*D*_eff_)*/P* (*D*_eff-isotonic_) of the measured probability of *D*_eff_ in different osmotic solutions, *P* (*D*_eff_), to that measured in isotonic solutions *P* (*D*_eff-isotonic_) for cells with either low or high acetylation. Total GEMs trajectories analyzed are n = 176140, 108348 and 155281 from n = 51, 45 and 40 cells for DMSO treated cells in either hypotonic, isotonic or hypertonic solutions, respectively, and n = 225775, 168719 and 170525 from n = 51, 47 and 39 cells for tubacin treated cells, from 3 independent experiments. (I) Image of a BEAS-2B cell expressing GFP-DCP1A-positive p-granules visualized by spinning disk confocal microscopy. Scale bar, 20 *µ*m. Inset: magnified view of p-granules. Scale bar, 2 *µ*m. (J) Measured MSD as a function of delay time *τ* for a representative p-granule trajectory as shown in the inset. Scale bar: 0.5 *µ*m. (K) Measured *D*_eff_ as a function of the apparent diameter of p-granules in DMSO- or tubacin-treated cells. Total p-granules trajectories analyzed are n = 9089 and 9183 from n = 45 and 46 cells from 3 independent experiments. (L) Measured MSDs for all the trajectories observed in each condition. The overall effective diffusion coefficient *D*_eff_ in each condition are 0.0098 ± 0.005 and 0.0059 ± 0.005 *µ*m^2^/s, respectively.

Cytoplasm fluidity is also governed by its viscoelastic properties (15, 39, 40, 62, 89). Previous studies have shown that increased acetylation can stiffen cells (28). To explore this, we exposed cells with varying levels of acetylation to osmotic environments ranging from hypoosmotic to hyperosmotic conditions (250–400 mOsm) (32, 38–40). This caused the cells to either swell or compress, allowing us to characterize GEM diffusivity in these altered cytoplasmic environments (Figs. 5E–H). In DMSO-treated cells, GEM diffusivity increased by 1.5-fold under swelling conditions and decreased by 52.4% under shrinkage conditions (Fig. 5F). However, in tubacin-treated cells, these changes were attenuated, with only a 1.3-fold increase under swelling and a 30.0% decrease under shrinkage (Fig. 5G). This modulation of GEM diffusivity was more apparent when the relative change in probability, *P* (*D*_eff_)*/P* (*D*_eff-isotonic_), was analyzed (Fig. 5H). These results indicate that hyperacetylated cells exhibit reduced water intake and loss when exposed to osmotic stress, modulating cytoplasmic viscoelasticity and rendering cells more resistant to osmotic pressure-induced volume changes.

We were next curious to examine whether microtubule acetylation affects the diffusive motion of larger, micron-sized particles in cells. We expressed GFP-tagged DCP1A, an mRNA-decapping co-activator and processing bodies marker, and tracked the motion of these micron-sized condensates in cells (89–92). DCP1A condensates varied greatly in size in BEAS-2B cells, and their motion was diffusive at short timescales (*<* 4 s) as the measured MSDs for individual trajectories were linear functions of delay time *τ* (Figs. 5I and J), consistent with previous findings (90, 91). Because the motion of DCP1A condensates is highly dependent on their sizes, we also computed their apparent diameters during tracking. We plotted the effective diffusion coefficient of individual condensates, *D*_eff_, as a function of their diameters in both DMSO and tubacin treated cells (Fig. 5K). The measured *D*_eff_ revealed a size-dependence with bigger condensates having smaller *D*_eff_ (Fig. 5K). Interestingly, tubacin treatment caused a 40.0% overall decrease of condensate diffusivity (Figs. 5K and L). Therefore, microtubule acetylation deceases the diffusive motion of micron-sized protein condensates in cells. In sum, we have demonstrated that microtubule hyper-acetylation stiffens the intracellular environment and slows down the diffusion of large particles in cells.

### Senescent cells display similar intracellular transport dynamics to hyperacetylated cells

We have demonstrated that microtubule acetylation is associated with cellular aging, and enhancing acetylation is correlated with decreased cytoplasm fluidity, which impacts vesicle transport, microtubule polymerization, and intracellular diffusion. We were therefore curious to determine if senescent cells display similar intracellular phenotypes to those observed in hyper-acetylated cells. Doxo-treatment decreased Rab6A vesicle mobile ratios by 16.4%, run time/total travel time ratios by 12.5%, peak run velocity by 20.0%, and run length and run time by 11.0% compared to controls (Figs. 6A-G), similar to what was observed in hyper-acetylated cells (Figs. 2M-Q). The effects on microtubule polymerization in senescent cells was even more pronounced. We measured 34.9%, 40.5%, 43.8% and 46.9% decreases in polymerization rate, polymerization length, polymerizing events and instantaneous velocity, respectively, and a 1.5-fold increase in pausing frequency in Doxo-treated cells compared to control cells (Figs. 6H-N). These findings suggest that the impaired intracellular transport and polymerizing dynamics likely originate from increased acetylation during cell senescence.

**Fig. 6.**
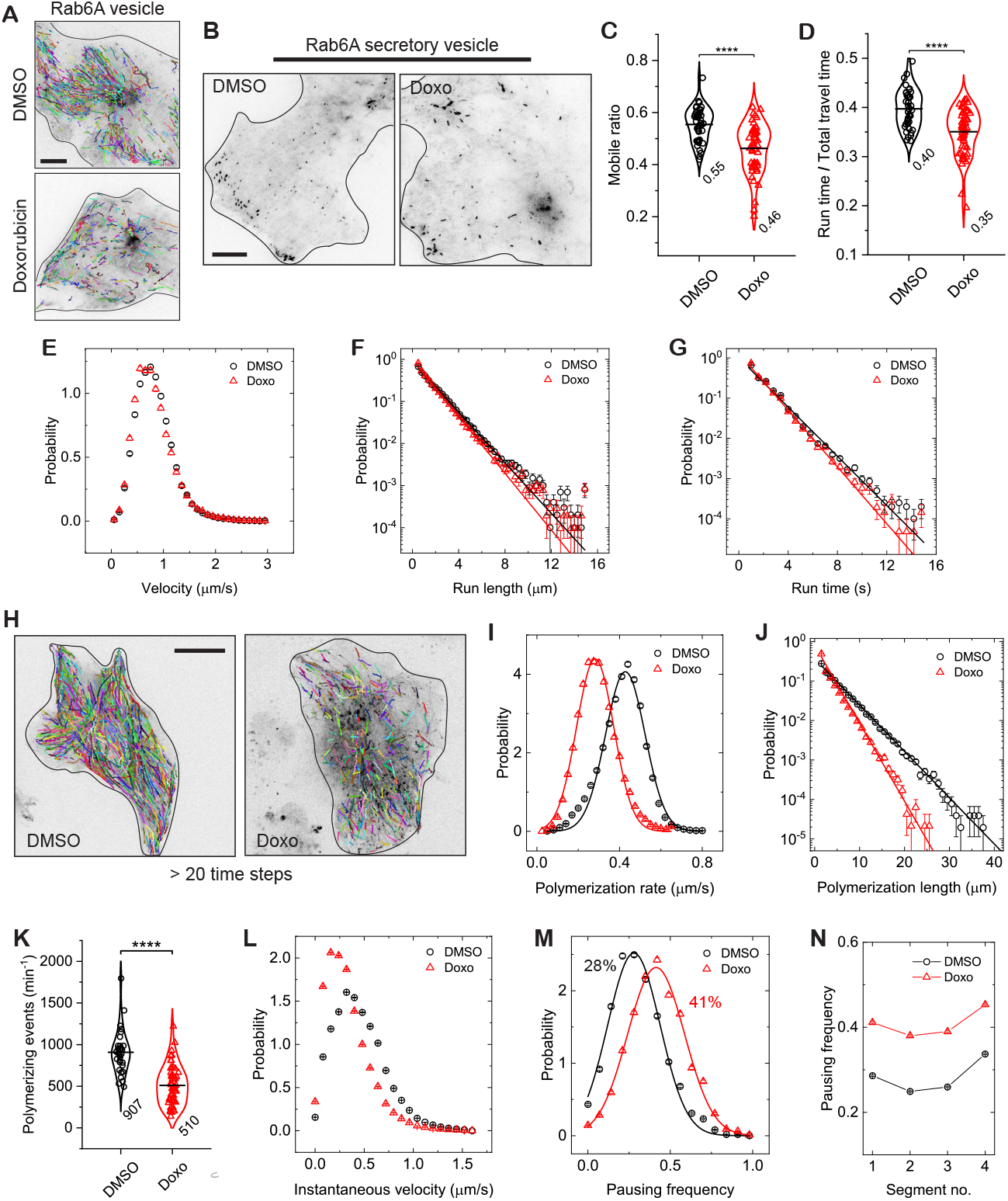
Cell senescence causes similar effects on anterograde transport and microtubule polymerization as hyperacetylation. (A) Representative trajectories of the mobile fraction of Rab6A-positive secretory vesicles for DMSO- and Doxorubicin-treated (Doxo) cells. Scale bar, 10 *µ*m. (B) Representative images of Rab6A vesicles in DMSO or Doxorubicin-treated BEAS-2B cells. Scale bar: 10 *µ*m. Insets show a magnified view of the vesicles in each condition. Scale bar: 1 *µ*m. (C and D) Measured mobile ratio and run time/total travel time of Rab6A-positive vesicles for DMSO- and Doxorubicin-treated cells. \*\*\*\**p <* 0.0001. Each data point represents the average of measured trajectories from a single cell. (E-G) Measured probability density functions (PDFs) of trajectory-based run velocity, *v*, run length, *L*_*p*_, and run time, *T*_*p*_, for Rab6A-vesicles in DMSO- and Doxorubicin-treated cells. The measured peak velocities are 0.75 ± 0.03 and 0.60 ± 0.03 *µ*m/s, respectively. The measured *L*_*p*_ and *T*_*p*_ both could be fitted to a simple exponential distribution (solid lines), with a characteristic *L*_*p*_ = 1.45 ± 0.02 and 1.29 ± 0.02 *µ*m, and *T*_*p*_ = 1.42 ± 0.02 and 1.23 ± 0.02 s, respectively. Total trajectories analyzed from the mobile fraction are n = 33127 and 34960 from n = 43 and 51 cells from 3 independent experiments. (H) Representative trajectories of EB1 comets in in DMSO or Doxorubicin treated cells. Scale bar, 20 *µ*m. (I and J) Measured PDFs of the polymerization rate (I) and the polymerization length (J) for individual EB1 comets in DMSO or Doxorubicin treated cells. The solid lines in (I) show the fitting of the data points to Gaussian distribution with the mean polymerization rates are 0.43 ± 0.01 and 0.28 ± 0.01 *µ*m/s, respectively. The measured polymerization length in (J) could be fitted to a simple exponential distribution (solid lines) with a characteristic length = 3.7 ± 0.15 and 2.2 ± 0.15 *µ*m, respectively, in DMSO or Doxorubicin treated cells. (K) Measured polymerizing events per minute in DMSO or Doxorubicin treated cells. \*\*\*\**p <* 0.0001. Total EB1 trajectories analyzed are n = 51509 and 47627 from n = 31 and 52 cells from 3 independent experiments. (L and M) Measured PDFs of the instantaneous velocity of EB1-comets and pausing frequency for individual EB1 comets in DMSO or Doxorubicin treated cells. The measured peak velocities are 0.32 ± 0.02 and 0.17 ± 0.02 *µ*m/s, respectively. The solid lines in (M) show the fitting of the data points to Gaussian distribution with the mean pausing frequencies are 28 ± 0.5% and 41 ± 0.2%, respectively. (N) Measured pausing frequency as a function of the trajectory-segment no. comets in DMSO or Doxorubicin treated cells.

## Discussion

Overall, our study establishes microtubule acetylation as a contributing factor to cytoplasmic mechanics and intracellular transport dysfunction during cell senescence and aging. By directly linking acetylation to phenotypes observed in senescent cells including decreased cytoplasmic fluidity, impaired vesicle motility, and disrupted microtubule dynamics, we provide mechanistic insight into how cytoskeletal modifications may drive age-related functional decline.

Aging is marked by declines in microtubule dynamics (20) and organelle transport (10, 17), yet the functional role of tubulin PTMs in these processes has remained enigmatic. We demonstrate that hyperacetylation reduces anterograde vesicle mobility and impairs microtubule polymerization through slower GTP-tubulin incorporation and dynamic pausing. These defects correlate with cytoplasmic density changes, evidenced by reduced diffusion of micron-sized condensates and resistance to osmotic stress. This aligns with studies showing microtubule stability modulates cytoplasmic viscoelasticity (28, 38) and suggests acetylation alters mechanical interactions between microtubules and cytoplasmic networks. Notably, acetylation’s effects are cargo-specific: Rab6A vesicles (predominantly anterogradely transport) are impaired, while Rab5 endosomes (predominantly retrogradely transport) remain unaffected, highlighting its selective impact on kinesin-driven processes.

Our findings reveal a reciprocal relationship: cytoplasmic viscosity modulates microtubule dynamics, while acetylation itself may lead to an increase in cytoplasmic viscoelasticity. This could create a self-reinforcing loop where age-related acetylation exacerbates transport defects, further reducing fluidity. For example, slowed microtubule polymerization in hyperacetylated cells may limit cytoskeletal remodeling, compounding cytoplasmic stagnation. Critically, the cytoplasmic changes we observe in hyperacetylated cells uniquely impacts large cargoes (e.g., Rab6A vesicles, DCP1A condensates, Figs. 2, 5), unlike metabolic defects, which broadly impair molecular motion (11, 13). This specificity positions acetylation as a potential mechanical rheostat, tuning fluidity to regulate transport efficiency. The resistance to osmotic stress further underscores its role in mechanical buffering, potentially synergizing with ion transporters (93) to maintain cell volume homeostasis.

Senescent cells phenocopy hyperacetylated cells, exhibiting reduced vesicle mobility, slower microtubule growth, and frequent pausing. These parallels, combined with elevated acetylation in aged *Drosophila*, position acetylation as a conserved hallmark of aging. Prior work linked acetylation to kinesin-1 recruitment (52, 59). Here, we observe elongated vesicles in hyperacetylated cells, which may suggest increased motor recruitment to overcome drag forces. However, persistent pausing implies compensatory mechanisms are insufficient in aging, leading to cumulative transport failure. This aligns with studies showing mitochondrial elongation and motility defects in aged neurons (10, 16) and supports the idea that acetylation-driven hardening exacerbates organelle dysfunction.

The exponential statistics of microtubule polymerization length suggest that termination of microtubule polymerization in cells occurs randomly in time and can be described by a Poisson process with a constant polymerization length despite the heterogeneous cellular environment (94). Elevation of acetylation, decrease of cytoplasm fluidity and drug-induced cellular senescence do not change the random nature of microtubule polymerization, but cause earlier terminations. This stability suggests that the local environment surrounding the growing microtubules is relatively homogeneous, facilitating a smoother pathway for microtubule polymerization and enhancing growth efficiency. This idea is further reinforced by the Gaussian statistics of microtubule polymerization rates (Figs. 3, 4, 6, S3 and S4). This pathway is perturbed by microtubule acetylation but not by detyrosination. Considering acetylation impedes the diffusive motion of micron-sized particles, we hypothesize that the polymerizing tips, through the interaction with a wide range of plus-ends associated proteins (80, 95, 96), may have an effective size comparable to micron-sized particles, and experience increased viscoelastic drag during polymerization upon increased acetylation.

We propose that microtubule acetylation acts as a mechanical aging clock, gradually impacting the cytoplasm to reduce fluidity and impair transport. This process may initiate early in aging, preceding metabolic decline or protein aggregation, and create a permissive environment for their progression. The conserved increase in acetylation across models (Fig. 1) and its sensitivity to osmotic cues (32) suggest it serves as a biomarker for cytoplasmic health. Restoring fluidity via acetylation modulation or motor activation (18, 97) could delay age-related decline, offering novel avenues for combating neurodegenerative and cardiovascular diseases linked to cytoskeletal dysfunction.

Our study bridges the gap between cytoskeletal biology and aging research, showing that microtubule acetylation is not merely a passive marker but an active driver of cellular aging. By impacting cytoplasm condensation and disrupting transport, acetylation creates a self-reinforcing cycle that contributes to senescence. Future work should explore how acetylation interacts with other aging hallmarks, such as protein aggregation (10) or metabolic decline (12). For instance, aggregated proteins could further condense the cytoplasm, synergizing with acetylation to impair transport. Additionally, the role of MAPs like MAP7 in sensing acetylation (32) warrants investigation, as their recruitment may mediate crosstalk between microtubules and cytoplasmic networks. Understanding these connections will clarify whether cytoskeletal dysfunction is a root cause or amplifier of aging, guiding therapies to promote healthy longevity.

## MATERIALS AND METHODS

### Cell lines

BEAS-2B cells (ATCC, CRL-9609) were maintained in Dulbecco’s modified Eagle’s medium (DMEM, Gibco), supplemented with 10% fetal bovine serum (FBS), 50 units/mL of penicillin and 50 µg/mL of streptomycin. All cell cultures were maintained in a 95% air/5% CO_2_ atmosphere at 37 ^°^C. The cell lines were routinely confirmed to test negative for mycoplasma contamination. For live cell imaging, cells were seeded at a density of approximately 1× 10^5^ cm^−2^ on a glass coverslip which was placed in a 35-mm polystyrene tissue-culture dish. Transfections were performed by using the FuGENE 6 (Promega) according to manufacturer’s instructions. Cells were generally transfected for 8 hours with 1 µg plasmids when the density reached ∼80% confluency.

### Fly stocks and husbandry

*Drosophila melanogaster* wild-type Oregon R stocks were raised on standard Bloomington cornmeal medium (Genesee Scientific 66-121). For aging experiments, all flies were maintained at 24 ^°^C under a 12-hr light:12-hr dark cycle. Equal numbers of male and female flies were collected the day of eclosion and housed together during aging. Experimental flies were transferred to fresh vials every 6-7 days.

### Antibodies

For Western blotting and immunostaining, we used rabbit monoclonal recombinant antibodies against *α*-tubulin with acetylation at *α*K40 (EPR16772, ab179484; Abcam), mouse monoclonal antibodies against *α*-tubulin with acetylation at *α*K40 (6-11B-1; Invitrogen), polyglutamylated tubulin (clone B3, T9822; Sigma) and *α*-tubulin (DM1A, T9026; Sigma), rabbit polyclonal antibodies against detyrosinated *α*-tubulin (AB3201; Sigma), chicken polyclonal antibodies against *α*-tubulin (ab89984; Abcam) and mouse monoclonal antibodies against BrdU (MoBU-1, Invitrogen). The following secondary antibodies were used: DyLight^*T M*^ 680 goat anti-mouse (35518, Invitrogen) and DyLight^*T M*^ 800 goat anti–rabbit (SA5-10036, Invitrogen) for Western blotting and Alexa Fluor 488–, 555–, and 647–conjugated goat antibodies against rabbit and mouse IgG (Invitrogen) for immunofluorescence.

### Generation of plasmids

The expression vectors used in this study were pCMV-*α*TAT1-EGFP (human isoform 4 modified from Addgene plasmid 27099), pCMV-*α*TAT1(D157N, catalytically dead mutant)-EGFP, pCMV-*α*TAT1-mScarlet, pCMV-*α*TAT1(D157N)-mScarlet, pEGFP-Rab6A (Addgene plasmid 49469), pEGFP-Rab5 (Addgene plasmid 49888), pCMV-VASH1-EGFP-P2A-SVBP, pCMV-VASH1(C168A, catalytically dead mutant)-EGFP-P2A-SVBP, pCMV-VASH1-mScarlet-P2A-SVBP, pCMV-VASH1(C168A, catalytically dead mutant)-mScarlet-P2A-SVBP, pDeltaCMV-VASH1-mScarlet-P2A-SVBP, pDeltaCMV-VASH1(C168A, catalytically dead mutant)-mScarlet-P2A-SVBP, EB1-tdTomato (Addgene plasmid 50825), pCDNA3.1-pCMV-PfV-GS-Sapphire (Addgene plasmid 116933; for mammalian expression of 40 nm-GEMs) and GFP-DCP1A (Addgene plasmid 153972).

VASH1 and SVBP proteins were cloned into a pDeltaCMV vector (a gift from Dr. Scott Hansen for expressing proteins at reduced levels) or pCMV vector split by a P2A selfcleaving peptide with a EGFP or mScarlet cassette at the C-terminus of VASH1 (32, 98). *α*TAT1 proteins were cloned into a pCMV vector with a C-terminal EGFP or mScarlet cassette. The single amino acid mutations were generated by PCR. All cloning were performed using Gibson assembly. All constructs were verified by DNA sequencing (Plasmidsaurus).

### Drug and osmotic treatments

To inhibit HDAC6 activity, cells were treated with tubacin (5 *µ*M, Sigma) in culture medium for 1 hr before being fixed or being imaged using TIRF or spinning-disk confocal microscopy. To induce cell senescence, cells were treated with Doxorubicin (200 ng/mL, Sigma), Camptocethin (CPT, 200 nM, Sigma) or Cisplatin (50 µM, Sigma) in culture medium for 24 hrs before being fixed or being imaged. To inhibit myosin II activity, cells were treated with Blebbistatin (10 *µ*M, Sigma) for 15 min before being imaged. The control groups were treated with DMSO. To manipulate the cytoplasm density, cells were treated with and imaged in extracellular osmotic environments ranging from hypoosmotic to hyperosmotic conditions (250–400 mOsm). These solutions were prepared by adding, respectively, 0.28 (∼400 mOsm, hypertonic), 0.2 (∼310 mOsm, isotonic control), 0.15 (∼250 mOsm, hypotonic) and 0.1 M (∼200 mOsm, hypotonic) mannitol to a hypotonic base solution (in millimolar: 40 NaCl, 5 KCl, 1 CaCl_2_, 2 MgCl_2_, 10 HEPES (pH 7.4), ∼91 mOsm) to maintain a constant ionic strength (32, 99). Cells were typically treated with the osmotic solutions 10 min before imaging.

### Western blotting

Flies were collected at the desired time point by flash freezing in liquid nitrogen and stored in −80 ^°^C. Groups consisted of 40 flies, including equal numbers of males and females. Heads and bodies were separated on dry ice by using frozen metal sieves (USA Standard Test Sieve No.25 710um, Newark Wire Cloth Company). For total protein level analysis, fly head and body were homogenized by using Kimble Pellet Pestle Motor and Pellet Pestle in fly lysis buffer containing 50 mM Tris-HCl, pH 8, 150 mM KCl, 2 mM MgCl_2_, 1 mM EGTA, 0.1% Triton X-100, 1 mM dithiothreitol (DTT), protease inhibitor (1:100, Roche) and phosphatase inhibitor (1:100, Sigma). The samples were heated in the Laemmli SDS sample buffer for 3 min at 95 ^°^C before electrophoresis. For electrophoresis separation, extracts were resolved in Mini-PROTEAN TGX Stain-Free protein gels (Biorad, cat 4568126) in standard Tris/glycine PAGE buffers and transferred to a nitrocellulose membrane (BioRad, cat 1620112) followed by blocking in 5% nonfat dried milk for 10 min at room temperature. All immunoblots were developed with corresponding primary antibodies (1:2000) and fluorescent secondary antibodies (1:4000), and visualized by infrared laser scanner (Odyssey CLx, Licor). Protein bands in Western blots were quantified using Fiji software (https://fiji.sc/).

### BrdU incorporation and staining

To assess the cell proliferation rate in senescent cells following Doxo, CPT and Cisplatin treatments, we measured the level of the thymidine analog BrdU (5-bromo-2’-deoxyuridine) following its incorporation into newly synthesized DNA by using anti-BrdU antibody. Briefly, drug-treated BEAS-2B cells were first incubated with 20 *µ*M BrdU at 37 ^°^C for 3 hrs to allow sufficient incorporation of BrdU. The cells were then rinsed with PBS for three times and fixed with 4% paraformaldehyde (PFA) and permeabilized with 0.2 M NH_4_Cl/PBS containing 0.5% Triton X-100 for 15 min each at room temperature. The cells were then incubated with 2 M HCl at 37 ^°^C for 30 min and then incubated with phosphate/citric acid buffer, pH 7.4 at room temperature for 10 min. The cells were then rinsed with PBS and blocked with 4% BSA in PBS for 2 hrs, which are ready for the incubation with primary and secondary antibodies.

### Immunostaining and confocal microscopy

For immunofluorescence cell staining, BEAS-2B cells were generally fixed in −20 ^°^C methanol for 10 min, and then blocked with 4% bovine serum albumin (BSA, Sigma) in PBS at room temperature for 2 hrs. Next, cells were incubated with primary antibodies against acetylated tubulin (1:200), detyrosinated tubulin (1:200), polyglutamylated tubulin (1:200) and *α*-tubulin (1:500), and corresponding secondary antibodies (1:200), each for 1 hr. For BrdU staining, an 1:100 dilution of primary antibodies was used. To stain F-actin and intermediate filaments, cells were fixed with 4% paraformaldehyde (PFA) and permeabilized with 0.2 M NH_4_Cl/PBS containing 0.2% Triton X-100 for 10 min each at room temperature, and then blocked with 4% BSA in PBS for 2 hours. F-actin was stained with Alexa-555-conjugated phalloidin (1:500, Invitrogen) for 1 hr. Intermediate filaments were labelled by incubating cells with a primary antibody against vimentin (1:200, Cell Signaling, 5741), and corresponding secondary antibodies (1:200), each for 1 hr. Cells were washed with PBS (5 min each for three times) before and after the incubation with secondary antibodies, and then mounted onto a microscope slide with ProLong^*T M*^ Gold Antifade Mountant (Invitrogen), and then examined by using spinning disk confocal microscopy. The spinning disk confocal was performed on an inverted research microscope Eclipse Ti2-E with the Perfect Focus System (Nikon), equipped with a Plan Apo 60× NA 1.40 oil objective, a Crest X-Light V3 spinning disk confocal head (Crest-Optics), a Celesta light engine (Lumencor) as the light source, a Prime 95B 25MM sCMOS camera (Teledyne Photometrics) and controlled by NIS elements AR software (Nikon). The fluorescent images for BEAS-2B cells were collected over a stack of vertical z-sections across the entire cell ∼4 *µ*m thickness. The final fluorescent images and their fluorescent intensities shown in the main text are based on the z-averaged images by using Fiji software (https://fiji.sc/).

For live-cell imaging of EB1-comets and DCP1A-positive P-bodies, a live-cell imaging chamber (H301-Nikon-TI-S-ER, Oko Labs) was equipped to the microscope to provide optimal culture conditions (95% air/5% CO_2_ atmosphere at 37 ° C) for cells during imaging. After cell transfection, the cell-containing glass coverslip was mounted on a coverslip holder (SC15012, Aireka Cells), which was finally mounted on the microscope. Images sequences for EB1-comets and DCP1A-positive P-bodies were typically recorded at 2.5 fps (frame per second) for 3 min and 5 fps for 3 min, respectively. The recorded images have 16 bits of gray scales and a spatial resolution of 700×700 pixels with the width of each pixel = 123 nm in our confocal optical setup.

### Total internal reflection fluorescence microscopy

TIRF microscopy experiments were performed on an inverted research microscope Eclipse Ti2-E with the Perfect Focus System (Nikon), equipped with a 1.49 NA 100× TIRF objective with the 1.5× tube lens setting, a Ti-S-E motorized stage, piezo Z-control (Physik Instrumente), LU-N4 laser units (Nikon) as the light source, an iXon DU897 cooled EMCCD camera (Andor) with an high-speed emission filter wheel (ET480/40M for mTurquoise2, ET525/50M for GFP, ET520/40M for YFP, and ET632/60M for mRuby2; Chroma). The microscope was controlled with NIS Elements software (Nikon). All live-cell experiments were performed in a live-cell imaging chamber (H301-Nikon-TI-S-ER, Oko Labs) that was equipped to the microscope to provide optimal culture conditions (95% air/5% CO_2_ atmosphere at 37 °C) for cells during imaging. After cell transfection, the cell-containing glass coverslip was mounted on a coverslip holder (SC15012, Aireka Cells), which was finally mounted on the microscope. Rab6A and Rab5 vesicles were typically recorded at 10 fps for 3 min. 40-nm-GEMs were recorded at 20 fps for 2 min. The recorded images have 16 bits of gray scales and a spatial resolution of 512×512 pixels with the width of each pixel = 107 nm in our TIRF optical setup.

### Single particle tracking and analysis

Single particle tracking (SPT) was performed using a homemade tracking program written in Matlab as previously described (32, 70), which is based on the standard tracking algorithm (100, 101). With this advanced SPT algorithm, we were able to obtain the position **r**(*t*) at time *t* for Rab6A-positive secretory vesicles, Rab5-positive early endosomes, EB1-comets, GEMs and DCP1A-positive P-bodies, and their trajectories were constructed from the consecutive images.

To characterize the mixed-motions of Rab6A-positive secretory vesicles and Rab5-positive early endosomes, we used an algorithm that could automatically identify and extract the states of diffusive diffusive “jiggling” movement and the states of directed “runs”, as described previously (32, 58, 68). Briefly, we determined the motion state of an arbitrary point in the trajectory by analyzing the turning angles around it at 5 different time scales. Trajectories containing no directed runs were considered as immobile. The mobile ratio was defined as the ratio of the number of mobile trajectories over the total number of trajectories in a single cell. The mobile ratio and run time/total travel time ratio generally characterize how active the motors are in moving vesicles from the “pauses” to the “runs” states at the whole cell level. Run velocity of each trajectory is obtained by dividing the total path length of all run segments to the total time spent in the run state. Run length and run time of each trajectory is obtained by averaging the path length or the dwell time of each run segments, respectively.

To characterize the intracellular dynamics of EB1-comets across the whole cell, we first selected the EB1 trajectories that exhibit clear directed motility (Fig. 3D). This is achieved by computing the mean squared displacements (MSDs), ⟨ Δ**r**^2^(*τ*)⟩ = ⟨ (**r**(*t* + *τ*) − **r**(*t*))^2^⟩, and fitted to ⟨ Δ**r**^2^(*τ*)⟩ = 4*D*_eff_*τ*^*α*^ to obtain the effective diffusion coefficient, *D*_eff_, and the scaling exponent, *α*. The *α* scaling exponent was used to classify EB1-comets motion as directed (*α* ≥ 1.5) (64, 102). Polymerization rate of individual EB1-comets was obtained by fitting the data to the parabolic function with ⟨Δ**r**^2^(*τ*)⟩ = *v*^2^*τ* ^2^. Polymerization length *L*_*p*_ is defined as *L*_*p*_ = *vT*_*p*_ with *T*_*p*_ is the lifetime of EB1-comets or polymerization time. Typically, *T*_*p*_ also features a simple exponential distribution. Polymerization events per minute is defined as the number of directed events detected in 1 minute in individual cells. Instantaneous velocity of EB1-comets *v*_inst_ is defined as *v*_inst_ = Δ*r/*Δ*t*, where Δ*r* is the path length that the EB1-comet travels over the shortest sample time (time stamp), Δ*t* = 0.4 s. The pausing frequency of individual EB1-comets was obtained by analyzing the turning angles, *θ*, during each time stamp. *θ* typically falls in the range −*π* ≤ *θ* ≤ *π*, with *θ* = 0 indicating directed forward movement and *θ* = −*π* or *π* indicating directed backward movement. Persistent directed motion or uninterrupted polymerization behavior is defined as −*π/*4 ≤ *θ* ≤ *π/*4, and with the rest defined as pausing behaviors. Pausing frequency is the ratio of pausing time to the total travel time of each trajectory. To quantify the temporal dependence of pausing frequency, individual trajectories (*>* 20 time steps) were equally cut into 4 segments as shown in Fig. 3I with segment 1 being the initiating phase and 4 being the ending phase of a polymerization event, respectively. The average pausing frequency of each segment was then computed. To characterize the apparent size of EB1 comets during microtubule polymerization, we used the *regionprops* function in Matlab to extract the shape properties of individual comets such as the short- and long-axis length, *a* and *b*, of the comet-shaped plus-ends, respectively. The apparent cap size, *L*, of individual trajectory is defined as the mean long-axis length, *b*. We also computed the average cap size in the pausing, *L*_pause_, and nonpausing states, *L*_nonpause_ of each trajectory, and compared their relative change, *L*_pause_*/L*_nonpause_, during polymerization.

To study the diffusion dynamics of GEMs, we first selected the mobile trajectories from the whole set of GEMs trajectories. This is achieved by computing the radius of gyration *R*_*g*_(*τ*) of each GEMs trajectory obtained over a time period of *τ*,

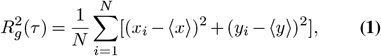

where *N* is the total number of time steps in each trajectory, *x*_*i*_ and *y*_*i*_ are the projection of the position of each trajectory step on the *x*− and *y*−axis, respectively, and ⟨*x*⟩ and ⟨*y*⟩ are their mean values. Physically, *R*_*g*_ quantifies the size of a GEMs trajectory generated during the time lapse *τ*. A cut-off value of 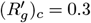 was used in the experiment, below which the GEMs trajectories are treated as immobile ones (32, 70). Here 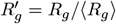 is the normalized radius of gyration, with ⟨*R*_*g*_⟩ being the mean value of *R*_*g*_. MSDs of individual mobile trajectories were then computed and fitted to ⟨Δ**r**^2^(*τ*)⟩ = 4*D*_eff_*τ* to obtain the trajectory-based effective diffusion coefficients of GEMs, *D*_eff_, in BEAS-2B cells.

The effective diffusion coefficients of DCP1A-positive P-bodies was obtained similar to that of GEMs. The apparent diameter of P-bodies was calculated by using the *regionprops* function in Matlab.

### Statistics

Data are expressed as mean ± s.e.m. unless specified otherwise. Graphs were created using Origin. Statistical tests were performed with two-tailed unpaired Student’s t-test. The statistical details of each experiment can be found in the figure legends.

## Acknowledgments

The authors wish to thank all the members of the Ori-McKenney and McKenney laboratories for their kind help and feedback. This work is supported by the NIH grant 1R35GM133688 to K.M.O.-M.

## Author Contributions

Y. S. and K. M. O. M. conceived the project and designed the experiments. R. M. and Y. S. performed the fly aging experiments. Y. S. performed all other experiments and analyzed the data. Y. S., R. M. and K. M. O. M. wrote the manuscript.

## Lead contact

Further information and requests for resources and reagents should be directed to and will be fulfilled by the lead contact, Kassandra M. Ori-McKenney.

## Materials availability

Cell lines and plasmids are available upon request from the authors.

## Data and code availability

Data generated and Matlab code used in this study are available upon request.

## Supplementary Videos and Figures

**Supplementary Video S1: EB1-comet dynamics in a BEAS-2B cell.** A 1 min video of the dynamics of growing microtubule plus-ends labelled by EB1-tdTomato in BEAS-2B cells (2.5 fps). EB1-tdTomato expressed in BEAS-2B cells exhibited a comet-like structure and directed motility from microtubule organizing center (MTOC) to the cell edge. Scale bar, 5 *µ*m.

**Supplementary Video S2: EB1-comet dynamics in a tubacin-treated BEAS-2B cell.** A 1 min video of the dynamics of growing microtubule plus-ends labelled by EB1-tdTomato in a tubacin-treated BEAS-2B cell (2.5 fps). Growing microtubule plus-ends displayed frequent pauses when they were pushing through the cytoplasm. Scale bar, 5 *µ*m.

**Fig. S1.**
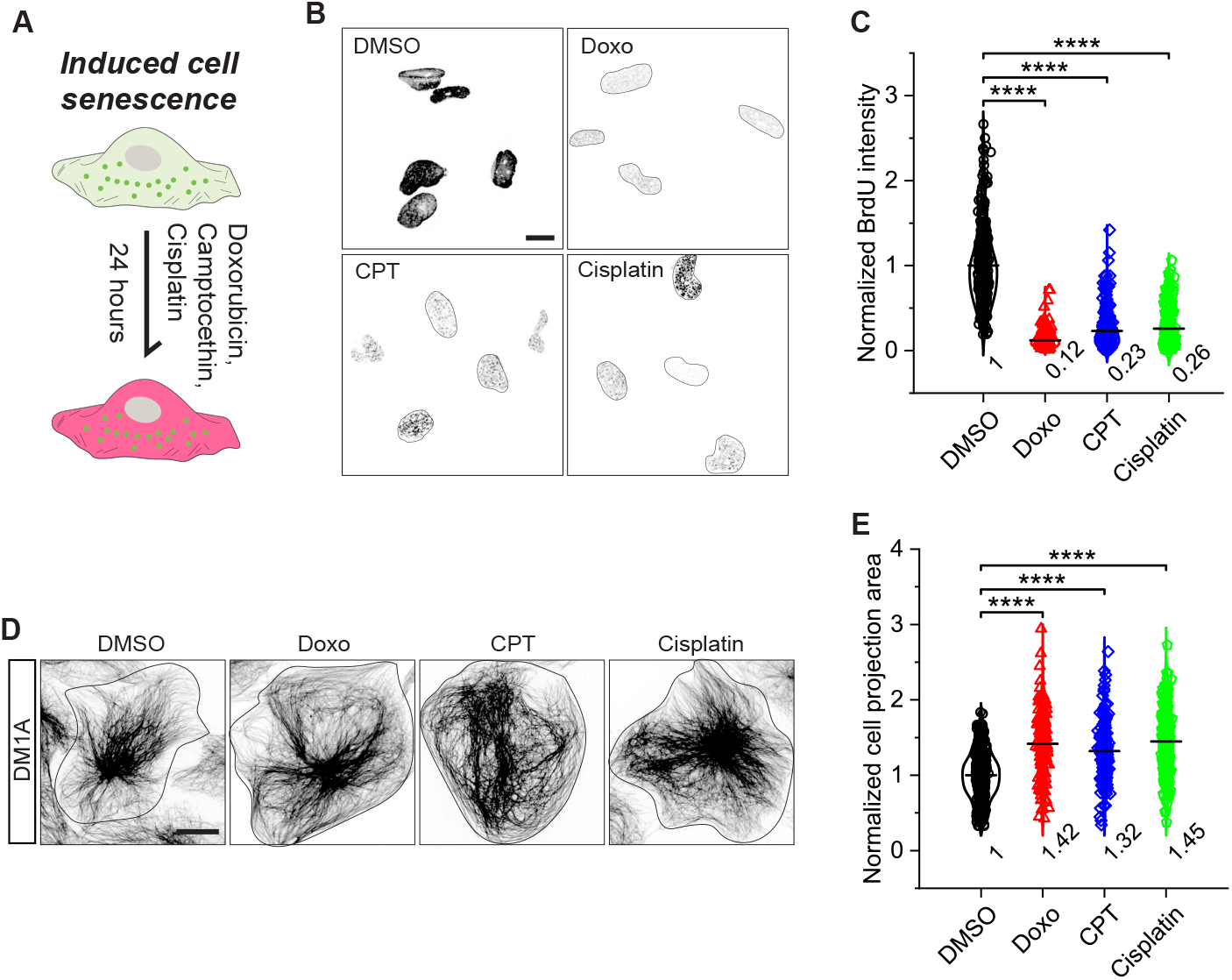
Doxorubicin, camptocethin and cisplatin treated cells display decreased DNA synthesis and increased cell projection area. (A) Schematic illustration of induced cell senescence by treating cells with Doxorubicin (Doxo), Camptocethin (CPT) or Cisplatin for 24 hours. (B and C) Representative images and quantification comparing BrdU incorporation into newly synthesized DNA in the nucleus for DMSO (control), Doxo (200 ng/mL), CPT (200 nM) or Cisplatin (50 *µ*M) treated cells. The black solid lines outline the nucleus shape. Scale bar: 20 *µ*m. n = 340, 347, 299 and 240 cells from 3 independent experiments, \*\*\*\**p <* 0.0001. (D and E) Representative images and quantification comparing cell projection area for cells treated with DMSO, Doxo, CPT or Cisplatin. Scale bar: 20 *µ*m. n = 459, 169, 172 and 176 cells from more than 3 independent experiments, \*\*\*\**p <* 0.0001.

**Fig. S2.**
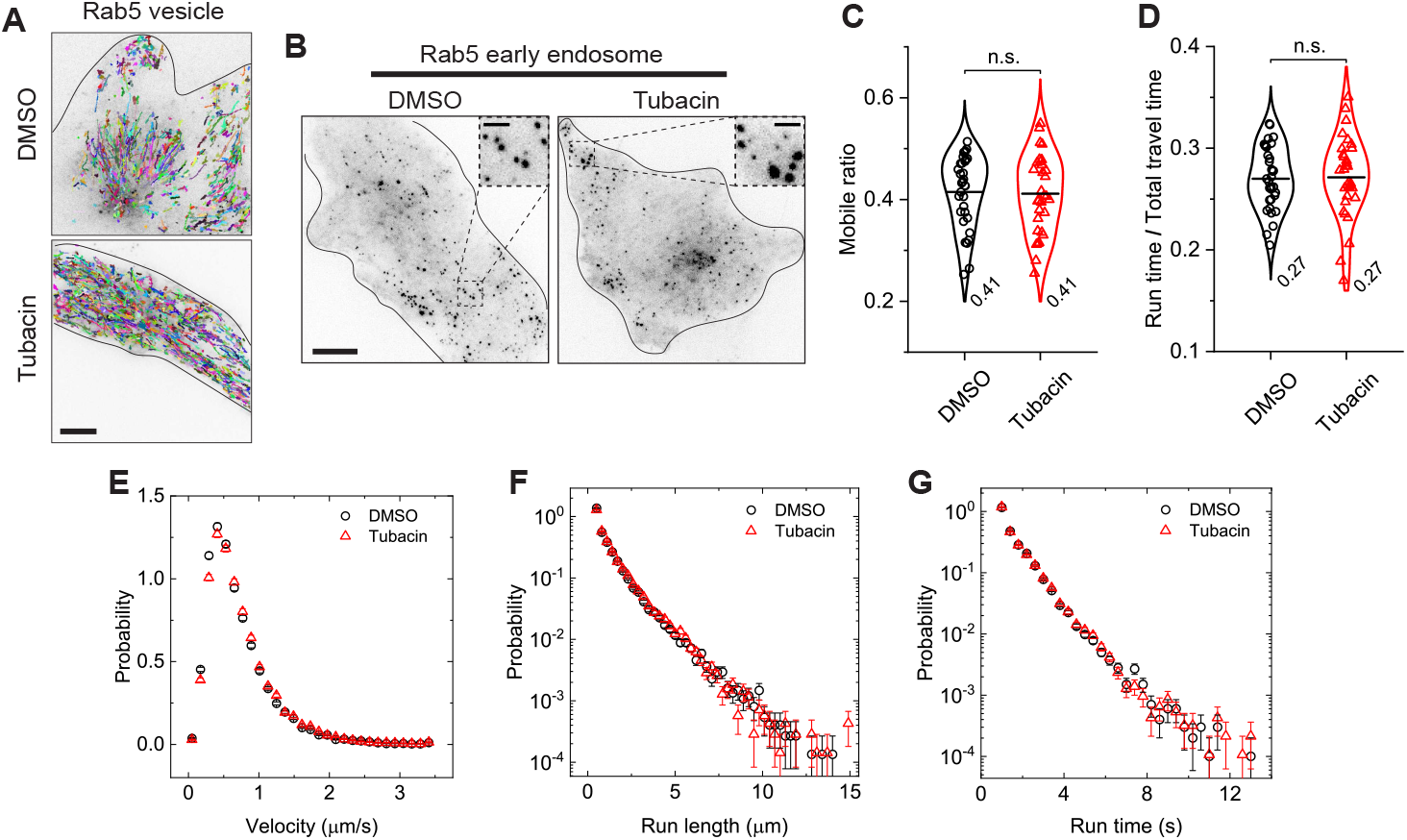
Microtubule acetylation does not affect retrograde transport. (A) Representative trajectories of the mobile fraction of Rab5-positive early endosomes for DMSO- and tubacin-treated cells. Scale bar, 10 *µ*m. (B) Representative images of Rab5 vesicles in DMSO or tubacin-treated BEAS-2B cells. Scale bar: 10 *µ*m. Insets show a magnified view of the vesicles in each condition. Scale bar: 1 *µ*m. (C and D) Measured mobile ratio and run time/total travel time ratio of Rab5 vesicles in DMSO or tubacin-treated cells. n.s. indicates *p >* 0.05. Each data point represents the average of measured trajectories from a single cell. (E-G) Measured probability density functions (PDFs) of trajectory-based run velocity, *v*, run length, *L*_*p*_, and run time, *T*_*p*_, for Rab5 vesicles in DMSO and tubacin-treated cells. The measured peak velocities for DMSO and tubacin treated cells are 0.41 ± 0.02 and 0.41 ± 0.02 *µ*m/s, respectively. The measured *L*_*p*_ and *T*_*p*_ both deviate from a simple exponential distribution with mean values *L*_*p*_ = 1.23 and 1.28 *µ*m, and *T*_*p*_ = 1.62 and 1.63 s, respectively, for DMSO or tubacin-treated cells. Total trajectories analyzed from the mobile fraction are n = 24921 and 23441 from n = 32 and 32 cells from more than 3 independent experiments. All violin graphs display all data points with means. *p*-values were calculated using a student’s t-test.

**Fig. S3.**
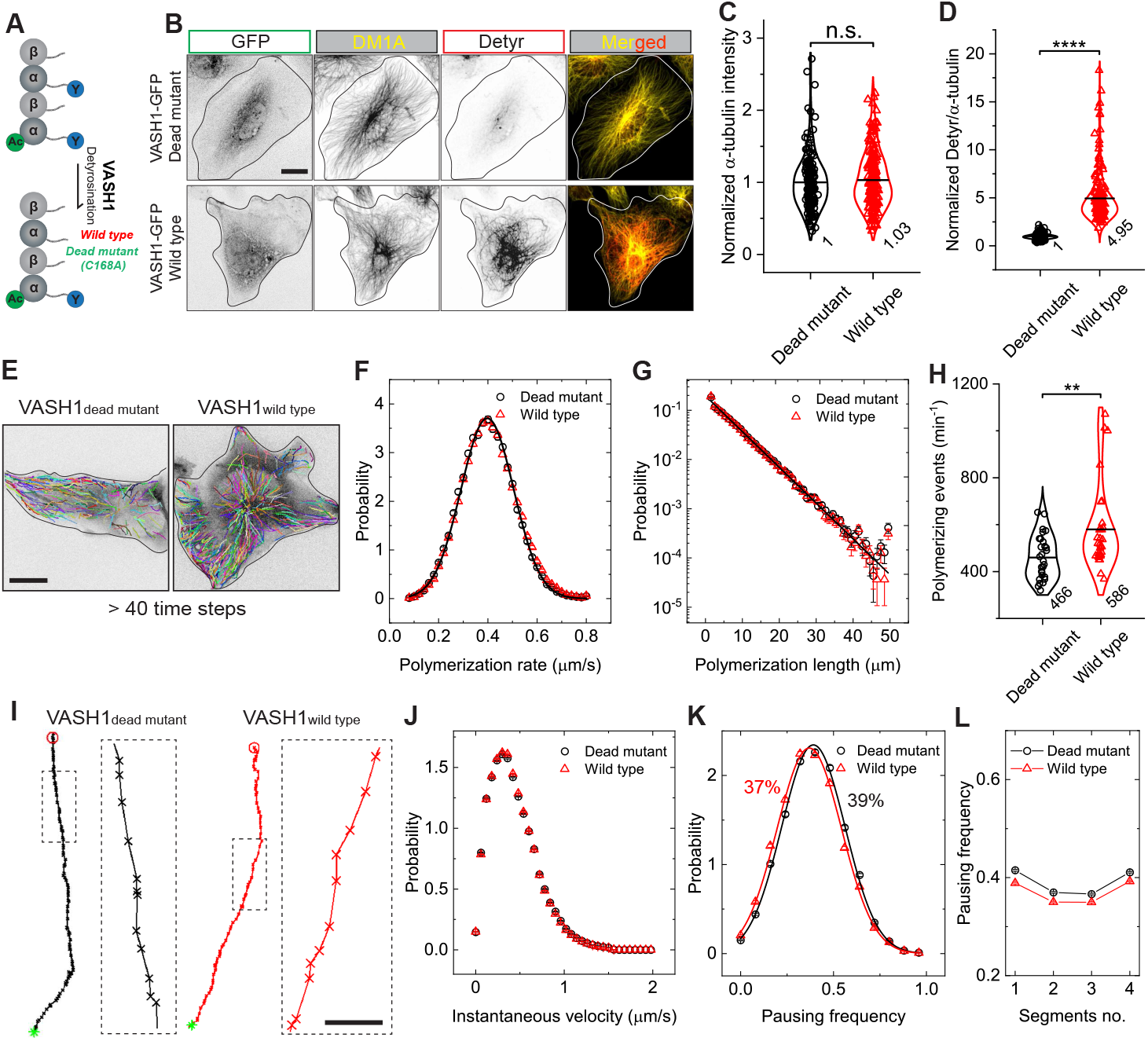
Microtubule detyrosination does not affect growing microtubule dynamics. (A-D) Representative images and quantification comparing *α*-tubulin intensity (C) and detyrosinated (Detyr) / total *α*-tubulin ratios (D) for cells transiently expressing the wild type (wild type) or catalytically dead mutant (C168A, dead mutant) of VASH1-EGFP-P2A-SVBP. n = 171 and 188 cells, respectively, from 3 independent experiments, \*\*\*\**p <* 0.0001. n.s. indicates *p >* 0.05. Scale bar: 20 *µ*m. (E) Representative trajectories of EB1 comets for cells transiently expressing the wild type or catalytically dead mutant of VASH1. Scale bar, 20 *µ*m. (F and G) Measured PDFs of the polymerization rate (F) and the polymerization length (G) for individual EB1 comets in cells transiently expressing the wild type (black circles) or the dead mutant (red triangles) of VASH1. The measured PDFs of polymerization rates largely overlap with each other and could be fitted to the Gaussian distribution (solid lines) with the mean polymerization rates = 0.40 ± 0.01 *µ*m/s. The measured PDFs of polymerization length also largely overlap with each other and could be fitted to a simple exponential distribution (solid lines) with a characteristic length = 6.0 ± 0.08 *µ*m. (H) Measured polymerizing events per minute in cells transiently expressing the wild type or catalytically dead mutant of VASH1. \*\**p* = 0.0022. Total EB1 trajectories analyzed are n = 54655 and 46349 from n = 30 and 29 cells from 2 independent experiments. (I) Magnified trajectory images showing EB1-comets movements in cells transiently expressing the wild type or catalytically dead mutant of VASH1. Scale bar, 0.5 *µ*m. (J and K) Measured PDFs of the instantaneous velocity of EB1-comets (J) and pausing frequency (K) in cells transiently expressing the wild type or catalytically dead mutant of VASH1. The measured peak velocities are 0.30 ± 0.05 and 0.31 ± 0.05 *µ*m/s, respectively. The solid lines in (K) show the fitting of the data points to Gaussian distribution with the mean pausing frequencies are 39 ± 0.2 % and 37 ± 0.3 %, respectively. (L) Measured pausing frequency as a function of the trajectory-segment no. in cells transiently expressing the wild type or catalytically dead mutant of VASH1. Total EB1 trajectories analyzed are n = 54655 and 46349 from n = 30 and 29 cells from 2 independent experiments. All violin graphs display all data points with means. *p*-values were calculated using a student’s t-test.

**Fig. S4.**
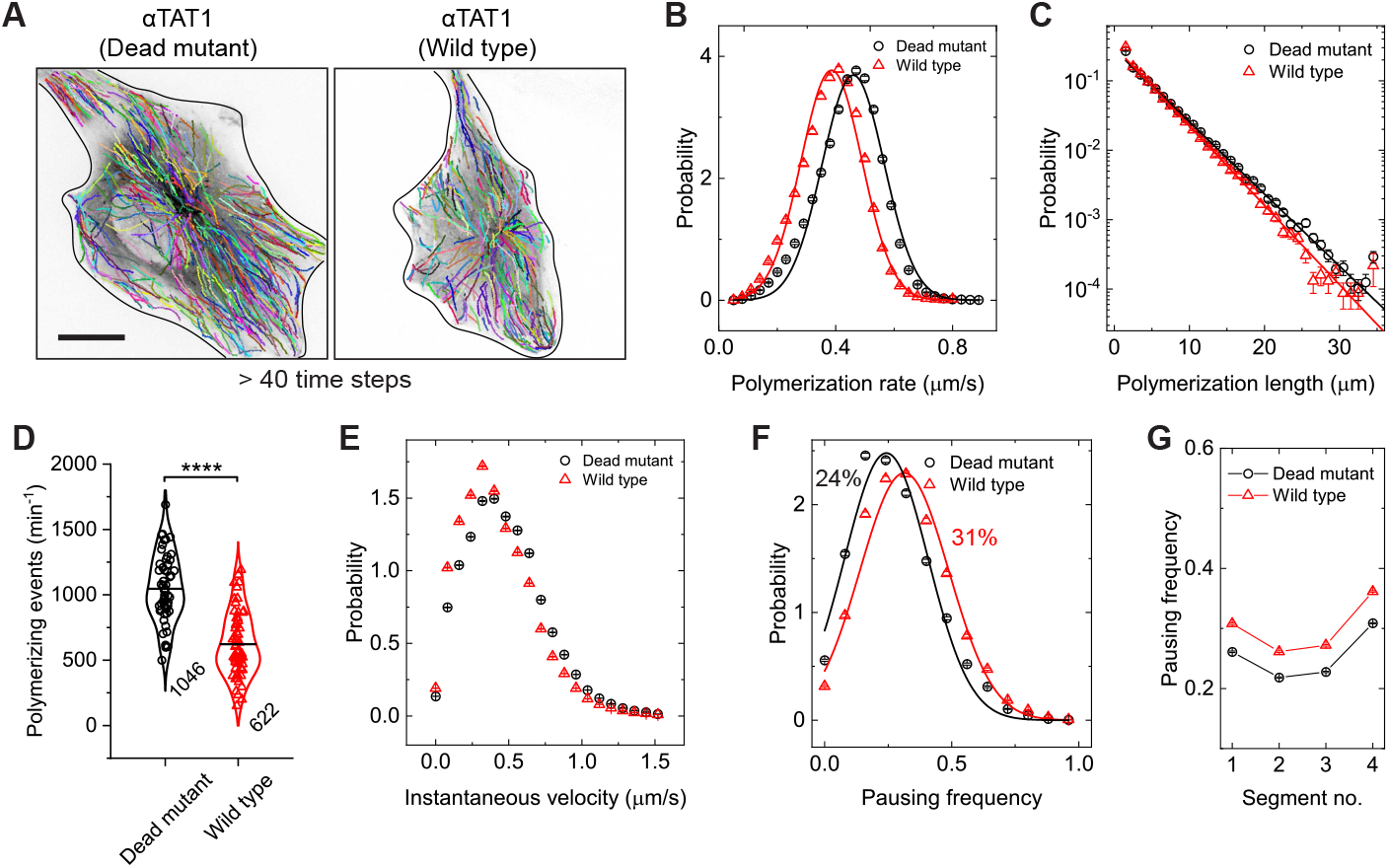
*α*TAT1 expression decreases growing microtubule polymerization rates and induces frequent pauses of growing microtubule plus ends. (A) Representative trajectories of EB1-comets in cells transiently expressing the wild type (wild type) or catalytically dead mutant (D157N, dead mutant) of *α*TAT1. Scale bar, 20 *µ*m. (B and C) Measured PDFs of the polymerization rate (B) and the polymerization length (C) for individual EB1-comets in cells transiently expressing the wild type or the dead mutant of *α*TAT1. The solid lines in (B) shows the fitting of the data points to Gaussian distribution with the mean polymerization rates are 0.38 ± 0.01 and 0.46 ± 0.01 *µ*m/s, respectively. The measured polymerization length in (C) could be fitted to a simple exponential distribution (solid lines) with a characteristic length = 3.80 ± 0.15 and 4.18 ± 0.15 *µ*m, respectively, for cells transiently expressing the wild type or the dead mutant of *α*TAT1. (D) Measured polymerizing events per minute in cells transiently expressing the wild type or the dead mutant of *α*TAT1. \*\*\*\**p <* 0.0001. Total EB1 trajectories analyzed are n = 88040 and 69099 from n = 54 and 63 cells from 3 independent experiments. (E and F) Measured PDFs of the instantaneous velocity of EB1-comets and pausing frequency for individual EB1 comets in cells transiently expressing the wild type or the dead mutant of *α*TAT1. The measured peak velocities in (E) are 0.30 ± 0.05 and 0.38 ± 0.05 *µ*m/s, respectively. The solid lines in (F) show the fitting of the data points to Gaussian distribution with the mean pausing frequencies are 31 ± 0.2% and 24 ± 0.5%, respectively. (G) Measured pausing frequency as a function of the trajectory-segment no. for cells transiently expressing the wild type or the dead mutant of *α*TAT1. All violin graphs display all data points with means. *p*-values were calculated using a student’s t-test.

**Fig. S5.**
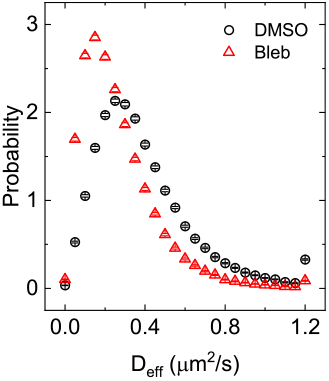
Blebbistatin treatment decreases GEMs diffusion. Measured PDFs of the effective diffusion coefficient *D*_eff_ for individual GEMs in DMSO or Blebbistatin treated cells. The measured peak *D*_eff_ are 0.27 ± 0.01 and 0.15 ± 0.01 *µ*m^2^/s, respectively. Total GEMs trajectories analyzed are n = 114319 and 79519 from n = 30 and 31 cells from 2 independent experiments.

